# Cell-type-specific encoding of prediction and reward in cortical microcircuits during novelty detection

**DOI:** 10.1101/2025.05.13.653877

**Authors:** Abdelrahman Sharafeldin, Hannah Choi

**Affiliations:** ML@GT, Georgia Institute of Technology, Atlanta, GA, USA; School of Mathematics, Georgia Institute of Technology, Atlanta, GA, USA

**Keywords:** Predictive coding, Efficient coding, Visual detection, Inhibitory interneurons, Disinhibitory circuit

## Abstract

Cortical circuits comprise diverse neuron types whose distinct activity patterns suggest specialized computational roles. Recent large-scale recordings reveal striking, cell-type-specific responses to novelty in cortex: excitatory neurons respond strongly to novel and unexpected stimuli, VIP interneurons respond more to novelty and omissions, while SST neurons are suppressed. What computational principles give rise to these dynamics? We introduce a normative model of a canonical cortical micro-circuit that jointly optimizes predictive coding, energy efficiency, and reinforcement learning under realistic connectivity constraints. By mapping algorithmic roles onto specific interneuron subtypes, the model reproduces absolute, contextual, and omission novelty effects observed experimentally. Critically, these emerge without hard-coded novelty detection and reveal a computational role for the VIP–SST disinhibitory motif in balancing representational capacity and metabolic cost. Mechanistic alternatives relying only on adaptation and Hebbian learning capture contextual but not absolute or omission effects. Our framework provides a unifying, falsifiable theory of how diverse cortical cell types implement coding principles underlying expectation and surprise.

## 1 Introduction

The brain operates in an environment that is both statistically structured and behaviorally dynamic. To efficiently perceive and act in the world, sensory systems must adapt not only to the physical properties of stimuli but also to the behavioral relevance of those stimuli in context. The efficient coding hypothesis ^1,2^ proposes that neural circuits are optimized to capture statistical regularities in sensory inputs and reduce redundancy given biological constraints such as metabolic cost. Predictive coding ^3,4^ builds upon this idea by proposing that the brain continuously predicts incoming sensory signals based on its internal representation of the world, computes the mismatch between predictions and incoming signals, and uses these prediction errors to update the internal model. This enables neural circuits to prioritize unexpected stimuli and suppress redundant, well-predicted inputs, conserving metabolic resources while enhancing sensitivity to behaviorally salient events.

While predictive coding offers a powerful normative theory, its biological implementation at the cortical circuit level remains an open question. Foundational work has shown how prediction and prediction-error units may be realized in cortical microcircuits ^3,5–8^, but these models often overlook local connectivity and abstract away the diversity and functional differentiation of inhibitory interneuron subtypes. They also do not consider how cortical processing unfolds in the context of behavior (although see Rao ^9^). Recent work has begun to reveal the diverse contributions of interneuron populations—particularly vasoactive intestinal peptide-expressing (VIP) and somatostatin-expressing (SST) cells—to shaping cortical computations ^10–12^. These subtypes participate in canonical circuit motifs such as the VIP-SST disinhibitory pathway, which enables dynamic, context-dependent gating of excitatory activity^13–15^ and is driven by reinforcement learning signals and reward-based feedback modulation ^16,17^. Despite increasing evidence for the involvement of such motifs in top-down modulation, attentional gating, and novelty processing ^18–22^, their role in predictive coding, particularly under behaviorally relevant conditions, remains poorly understood.

Here we address these limitations by introducing a biologically grounded *normative* model of cortical microcircuits that unifies predictive coding, energy efficiency, and reinforcement learning. Unlike earlier predictive coding frameworks that abstract away interneuron diversity or ignore behavioral context, our approach assigns distinct computational roles to excitatory and inhibitory subtypes in a connectivity-constrained microcircuit. SST neurons encode the mean of top-down predictions, VIP neurons capture their uncertainty, PV neurons represent the precision of feedforward drive, and excitatory neurons are divided into representational and prediction-error subpopulations. This circuit is further constrained by Dale’s law–the principle that each neuron releases either excitatory or inhibitory neurotransmitters. By optimizing multiple objectives during a naturalistic behavioral task, the model links coding principles directly to the observed diversity of cell-type-specific responses. In doing so, it bridges the gap between abstract normative theories and mechanistic circuit models, providing a framework in which hallmark interneuron motifs—such as VIP-SST disinhibition—emerge as solutions to competing computational demands.

We evaluate our model in a recent novelty-detection paradigm used by Garrett et al. ^23^. Novelty detection is a key behavioral context that can be especially understood through the lens of predictive coding. Ethologically, detecting novel stimuli, such as threats, opportunities, or salient changes in the environment, is fundamental for an animal’s survival. Indeed, neural responses to novelty have been recorded across multiple brain areas and species ^24–30^. Importantly, responses to three types of novelty have been studied: *absolute* novelty, which distinguishes between previously observed (familiar) and unobserved (novel) stimuli ^31,32^, *contextual* novelty, where an otherwise familiar stimulus appears in an unexpected context ^33,34^, and *omission* novelty, where an anticipated stimulus fails to appear ^35^. These distinct forms of surprise all reflect violations of internal expectations. Several computational studies have explored predictive coding as a framework for novelty detection^36–38^, but these typically focus on abstract architectures that are not based on cortical microcircuit structure and do not aim to replicate experimentally observed cell-type-specific responses.

Garrett et al. ^23^ recorded large-scale neuronal responses in the visual cortex of mice performing a visual change detection task. Using two-photon calcium imaging in excitatory, VIP, and SST neurons across the primary and secondary visual areas, they measured activity during repeated presentations of natural images, interspersed with image changes and unexpected omissions. Their data revealed strikingly robust cell-type-specific novelty responses: VIP neurons responded strongly to novel stimuli, image changes, and omissions; SST neurons were suppressed by novel stimuli; and excitatory neurons showed enhanced responses to novel stimuli and image changes. The matching effects in VIP and excitatory neurons are thought to reflect disinhibition, where VIP activity suppresses SST neurons, thereby releasing excitatory neurons from inhibition ^13,23,39,40^. These findings suggest distinct computational roles for each population in encoding sensory predictions, uncertainty, and errors—yet the underlying circuit mechanisms remain unclear.

We show that our model, trained on the naturalistic change detection in Garrett et al. ^23^, reproduces the diversity of novelty effects observed experimentally, including absolute, contextual, and omission responses, across all three cell types. Crucially, these effects emerge naturally as a consequence of optimizing predictive coding, energy efficiency, and reinforcement learning objectives using a biologically-constrained network of functionally specialized subpopulations. This places our work in the tradition of normative modeling studies such as Olshausen and Field ^41^, Rao and Ballard ^42^, and Yamins et al. ^43^, where optimizing computational principles in realistic tasks led to the spontaneous emergence of hallmark biological properties. Our results also support the hypothesis that novelty responses emerge naturally in networks that learn efficient neural representations ^38,44,45^, in contrast to models that require dedicated circuits for novelty detection ^46,47^ and uncertainty-based responses ^48^.

By conducting ablation analyses on different parts of the model, we demonstrate that predictive coding is necessary for generating absolute novelty responses, while reinforcement learning and reward modulation produce contextual novelty effects in VIP neurons. These analyses further reveal a critical role for VIP-SST disinhibitory motif in balancing representational capacity with energy efficiency. Finally, we tested an alternative model based on response adaptation and Hebbian learning. While biologically plausible, the alternative model only captured contextual novelty and failed to produce the absolute novelty effects seen experimentally, suggesting that mechanistic principles alone are insufficient and that assigning computational roles to specific cell types is essential for explaining the full range of novelty responses. Our framework thus provides a falsifiable, general account of how canonical cortical motifs implement predictive and efficient coding principles, unifying data-driven observations of novelty responses with a theory of cell-type-specific computation.

## 2 Results

### Task Setup

We train the model on a change detection task similar to the one described in Garrett et al. ^23^. A set of images is used to construct change sequences, where one image is presented repeatedly at regular times until a change occurs and a different image is presented for the remainder of the sequence. Each image is shown for 2 time steps and followed by 4 time steps of blank stimulation (zero input), as shown in Figure 1E. During training, sequences are constructed from a set of familiar images, while during testing, sequences are built from a novel image set. Both sets consist of natural scene images, examples of which are shown in Figure 1B. The task also includes omission trials where an expected image presentation is omitted and replaced by a blank stimulus at a random time in the sequence (Figure 1E, bottom). During testing, each sequence has a 30% chance of containing an omission. No omission trials occur during training, consistent with the experimental paradigm. See Methods for more details on sequence construction, image datasets, and the experimental change detection task.

**Figure 1:**
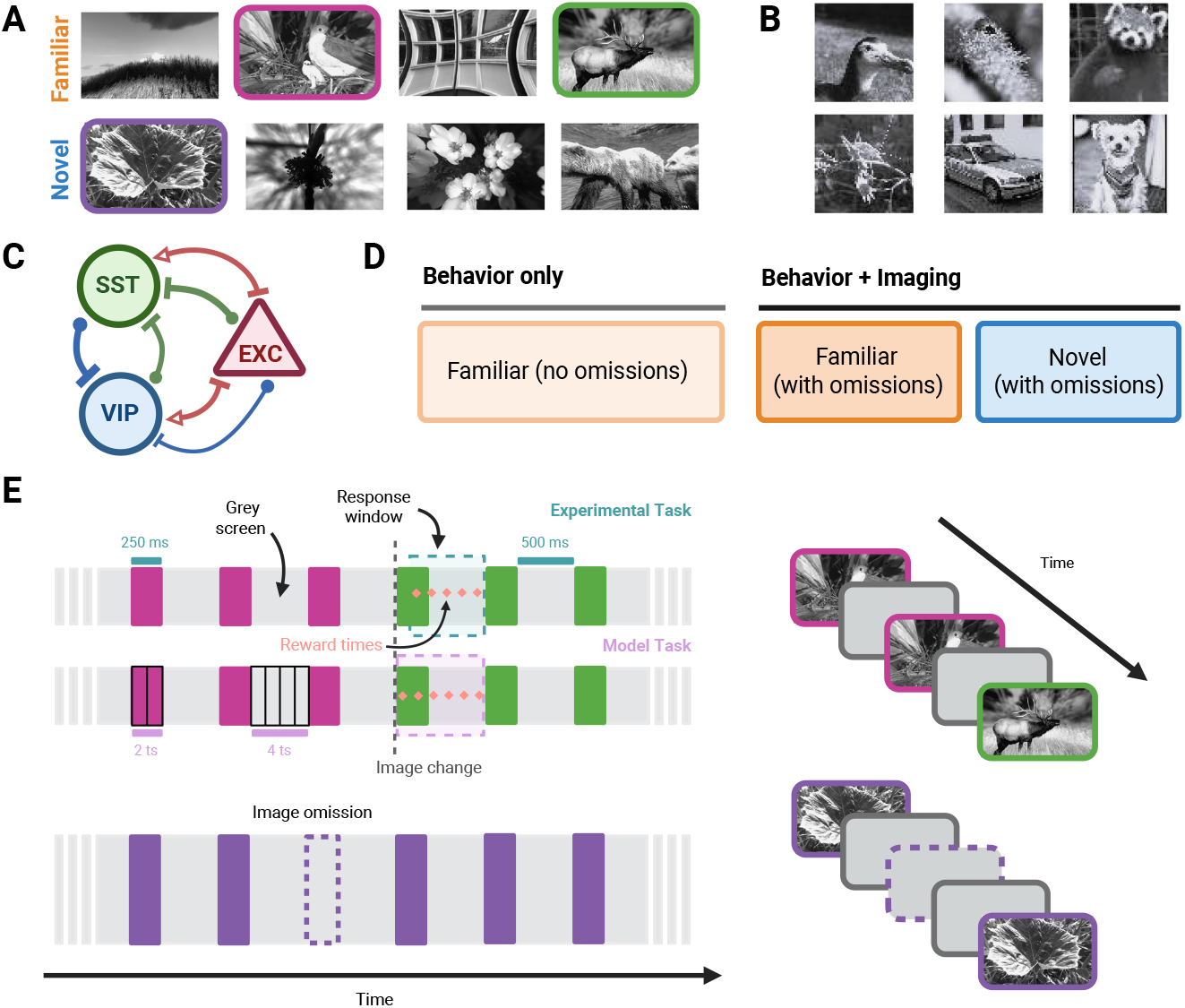
Experimental and model task setup. (A) Example images from the familiar and novel image sets used in the experimental change detection task (reproduced with permission from Garrett et al. ^23^, data available from the Allen Institute’s Visual Behavior (2P) dataset at https://portal.brain-map.org/circuits-behavior/visual-behavior-2p). (B) Example images used in the model change detection task. These images are a subset of the TinyImageNet dataset ^49^ converted to grayscale. (C) Schematic of part of the mouse visual cortical circuit containing the neuron populations recorded in experiment. (D) Session schedule for behavioral training and calcium imaging used in the experimental setup. Mice were first trained on familiar image change sequences without imaging. Then, neurons were recorded during familiar and novel sessions with omission trials. (E) Top: sample image change sequence in the experimental and model tasks. Different-colored squares correspond to different image presentations. Bottom: sample image sequence for an omission trial, where an expected image presentation is skipped and replaced by a gray screen.

In the experimental task, mice are trained to lick for a water reward within a response window after detecting a change (Figure 1E). Similarly, our model selects at each time point one of two actions: lick or no lick. For each lick in a 6-time-step window following the change, it receives a reward of 10 units, minus a 2-unit cost for licking, for a net of +8 units per lick. Outside of this response window (i.e., when no change has occurred), a lick yields a reward of − 2 units (no water reward but still incurring the action cost). Not licking always yields a reward of 0 units. This structure encourages the model to reliably detect the change while penalizing false alarms.

### Cortical circuit model

Our model consists of four hierarchical layers designed to mimic the different stages of cortical processing in the brain. These layers are the input layer (Layer 0), the intermediate layer (Layer 1), the sensory representation layer (Layer 2), and the higher-level processing layer (Layer 3). The architecture and connectivity of the model are described below and depicted in Figure 2.

**Figure 2:**
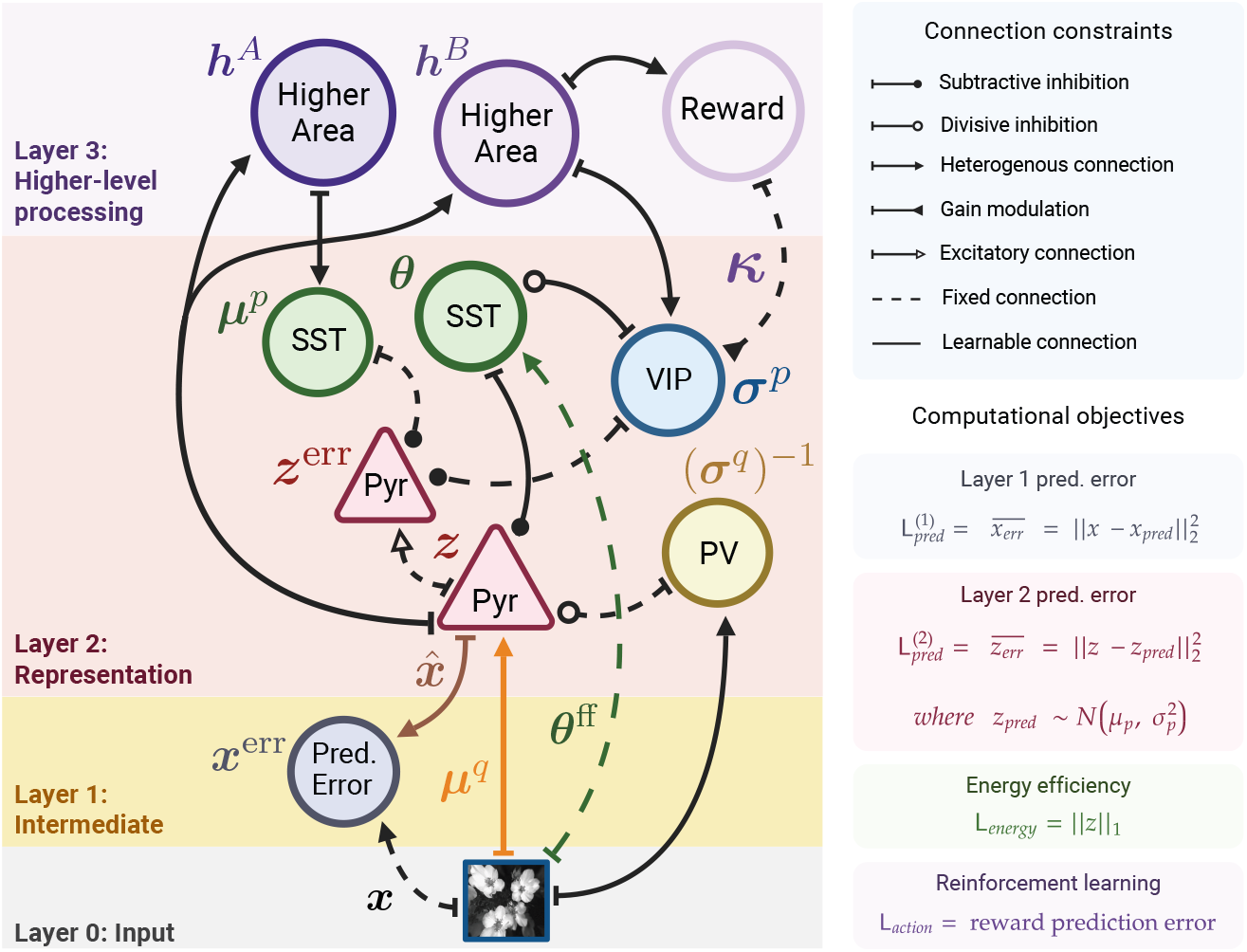
Model architecture, connectivity, and computational objectives. Schematic of the cortical circuit model we develop in this work. All solid connections are learnable and parametrized by connection weight matrices, while dashed connections are fixed and determined by their role in the algorithmic implementation (see Methods). All intra-layer connections are constrained to obey Dale’s law, while inter-layer connections can have both excitatory and inhibitory weights. Connectivity in Layer 2 was determined based on experimental evidence of interneuron connectivity in real cortical circuits (see Methods). Computational objectives (bottom right) combine to form the loss function used to train the model. The notations 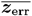 and 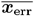 indicate population averages.

#### Input layer

The input layer simulates the retina in the visual pathway and is where the stimulus presentation occurs. At this layer, feedforward connections from the input stimulus project to Layer 1 prediction error population (*x*^err^) as well as to excitatory pyramidal (Pyr), inhibitory SST, and inhibitory PV subpopulations in Layer 2.

#### Intermediate layer

The intermediate layer corresponds to the intermediate stages of the visual processing pathway, such as the lateral geniculate nucleus (LGN) and other thalamic regions. It also captures the role of layer 4 in primary visual area, which is the main cortical recipient of thalamic input. At this layer, prediction error neurons *x*^err^ compute the difference between the input *x* and its reconstruction 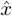 generated by feedback from the excitatory *z* neurons in Layer 2.

#### Representation layer

The sensory representation layer models early visual cortical areas–the primary (VISp) and secondary (VISl) areas of the mouse visual cortex, where experimental data was recorded ^23^. Functionally, it aligns with layer 2/3 in the laminar organization, which integrates input from earlier stages and participates in intracortical processing and feedforward projection to higher areas. It contains four main populations: excitatory (Pyr), SST, VIP, and PV neurons. The excitatory population is divided into representation (*z*) and prediction error (*z*^err^) subpopulations, motivated by experimental evidence for such subpopulations ^28,50–53^. Similarly, SST neurons consist of two subpopulations: SST (*θ*) and SST (*µ*^*p*^). Excitatory *z* neurons receive feedforward connections from the input layer and are inhibited by PV and SST (*θ*) neurons. They send feedforward projections to higher areas in the higher-level processing layer. Prediction error neurons (*z*^err^) receive input from the excitatory *z*, VIP, and SST (*µ*^*p*^) populations. VIP neurons receive feedback from the higher-level processing layer and is modulated by top-down reward signals. They inhibit SST (*θ*) neurons, forming a disinhibitory pathway. SST (*θ*) neurons receive weak feedforward input and inhibit the excitatory (*z*) neurons. SST (*µ*^*p*^) neurons receive feedback from the higher-level processing layer and inhibit the excitatory prediction error (*z*^err^) neurons. Finally, PV neurons receive feedforward input and provide divisive inhibition to the excitatory representation (*z*) neurons.

#### Higher-level processing layer

The higher-level processing layer represents cortical areas involved in recurrent processing of input sequences and reward-based modulation. It consists of Higher Area A, Higher Area B, and the Reward area. Higher Areas A and B receive feedforward input from the excitatory *z* population in the representation layer. The Reward area receives input from Higher Area B, and sends a modulatory feedback signal to the VIP population in the representation layer. Neurons *within* each of these three areas are recurrently connected.

All intra-layer connections in the model, whether learned or fixed, obey Dale’s law and are constrained to be either excitatory or inhibitory. Inter-layer connections are not subject to this constraint and may be of either type. This is because, biologically, long-range inter-layer connections may go through intermediate steps that result in a mixed excitatory/inhibitory net effect at target regions. Further details on the model architecture, Dale’s law constraints, synaptic weight matrices, and algorithmic implementation are provided in the Methods section.

### Computational objectives

The central hypothesis of this work is that distinct novelty responses in excitatory, VIP, and SST populations can be explained by mapping these cell types onto algorithmic nodes within an energy-efficient predictive coding framework that also incorporates reinforcement learning. Below, we outline the computational objectives of the model—predictive coding, energy efficiency, and reinforcement learning. Then, we describe how the algorithmic components of these objectives map onto specific interneuron subtypes.

In the predictive coding framework, the brain maintains a generative model of the world where latent states *z* give rise to observations *x*. This generative model consists of a likelihood distribution *p*(*x* | *z*) and a prior distribution *p*(*z*), which encodes expectations about the latent states given historical data. The parameters of this generative model can be learned by maximizing the log evidence, given by

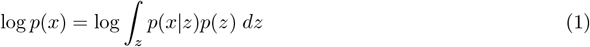

However, since directly maximizing log *p*(*x*) is intractable, we maximize a variational lower bound obtained by introducing a variational posterior *q*(*z* | *x*) which approximates the true posterior *p*(*z* | *x*). This variational bound is commonly referred to as the Evidence Lower Bound (ELBO) and is given by

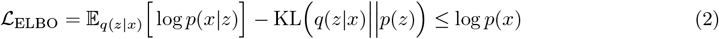

The first term in the ELBO encourages the model to reconstruct observations faithfully. When parameterizing *p*(*x* | *z*) as a Gaussian distribution with mean 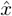 and identity covariance, this first term becomes (negative) the mean squared error (MSE) between the observation *x* and the reconstruction (or prediction) 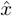. The second term in the ELBO regularizes the approximate posterior to stay close to the prior. However, since neural circuits are unlikely to compute the exact KL divergence between distributions, we perform this regularization implicitly in a biologically plausible manner by minimizing the MSE between the prior’s predicted state, 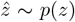, and the posterior’s inferred state, *z* ∼ *q*(*z* | *x*). Using this approach, the predictive coding component of our model becomes similar to the biological implementation proposed by Rao and Ballard ^3^, where higher brain areas relay predictions to lower areas through feedback connections and the prediction error at each layer is minimized. This yields the following two prediction error loss functions (one for each of Layer 1 and Layer 2)

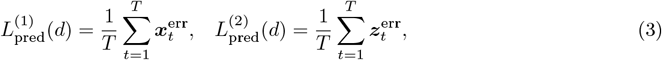

where *d* is a given image sequence, *T* is the duration of that sequence (in time steps), and 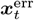 and 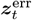 are the activities of the prediction error neurons at Layer 1 and Layer 2, respectively.

Besides minimizing prediction errors for perceptual inference, the model must also learn to take actions that maximize reward in the behavioral task. This is achieved using an actor-critic reinforcement learning framework, in which the critic estimates expected rewards and the actor selects actions to maximize them. Learning is driven by reward prediction errors (RPEs), computed as the discrepancy between expected and received rewards. The model updates its parameters to minimize these RPEs over time, aligning with dopaminergic reinforcement learning mechanisms observed in biological systems ^54,55^. Minimizing RPEs also fits well within our framework, which is centered around predictive processing. This computational goal yields the following behavioral loss function for a given image sequence *d*

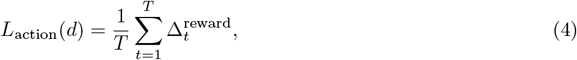

where 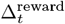 is the RPE at time step *t*, computed as the difference between actual and predicted rewards at that time step (see the Algorithmic Implementation section in Methods).

The energy efficiency component of our model encourages sparse neural representations by minimizing the *l*_1_-norm of the inferred latent state *z* which is represented by the excitatory representation neurons. This promotes sparse coding, where only a subset of neurons are active at any given time, reducing metabolic costs while maintaining representational efficiency ^2^. By balancing sparsity with prediction accuracy, the model avoids unnecessary redundancy in neural activity, allowing it to represent information more efficiently. The energy efficiency loss function for a sequence *d* is given by

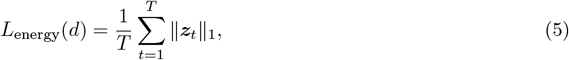

where ***z***_*t*_ is the activity of the excitatory *z* neurons at time step *t*.

The final loss function used to train the model is a combination of the above objectives, and is given by

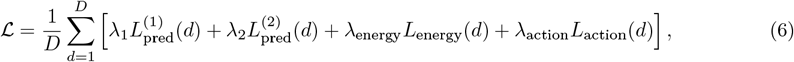

where *D* is the number of sequences in the training dataset, and *λ*_1_, *λ*_2_, *λ*_energy_, and *λ*_action_ are scalar *weights* that balance the different objectives’ contribution to the final loss. In practice, these weights are annealed throughout training to guide optimization (see Methods).

### Correspondence between algorithmic and biological components

Below, we outline the proposed mapping between specific interneuron populations and the algorithmic components of the model. This mapping is informed by novelty responses and known functional properties of excitatory, VIP, and SST neurons, for which experimental data are available. As PV neurons were not recorded experimentally, their assigned role is based on known connectivity and function, and the resulting novelty responses constitute model predictions to be tested experimentally. For further justification and algorithmic implementation details, see Methods.

#### PV neurons

PV neurons receive feedforward drive from the input layer and represent the *precision*, (*σ*^*q*^)^−1^, of the posterior distribution *q*(*z*|*x*).

#### VIP neurons

VIP neurons receive feedback input from higher areas and encode the uncertainty of top-down predictions by representing the standard deviation, *σ*^*p*^, of the prior distribution *p*(***z***).

#### SST neurons

There are two subpopulations of SST neurons in our model: SST (*µ*^*p*^) and SST (*θ*). The SST (*µ*^*p*^) population represents the mean, *µ*^*p*^, of the prior distribution *p*(*z*). The SST (*θ*) population serves as part of the VIP-SST disinhibitory pathway and acts as the primary source of inhibition onto excitatory neurons.

#### Excitatory neurons

The excitatory population in our model consists of two subpopulations: representation neurons (*z*) and prediction error neurons (*z*^err^). Representation neurons receive feedforward drive from the input layer, representing the mean, *µ*^*q*^, of the posterior distribution *q*(*z* | *x*). This feedforward drive, combined with PV input, effectively produces a sample from the posterior distribution, modified by inhibitory input from SST (*θ*) neurons. Prediction error neurons encode the mismatch between top-down predictions and bottom-up inputs at the representation layer, computed through local interactions with excitatory and inhibitory neurons receiving feedforward and feedback input, respectively.

### Training, internal predictions, and sequence responses

We trained 32 instances of our model, each corresponding to a different random seed. For each instance, we randomly draw a set of 8 *familiar* images and a set of 8 *novel* images from the image dataset. The first set is used to construct the image change sequences used for training, while the second set is used to construct test sequences. The model is trained for 70 epochs with backpropagation to minimize the objective function in Eq. 6. We set *λ*_1_ = 1, while the weights *λ*_2_, *λ*_energy_, and *λ*_action_ are annealed throughout training according to the schedules in Eq. 37.

The training trajectories of the various loss terms, along with their annealing schedules, are shown in Figure 3A–F. At the start of training, the model rapidly minimizes the Layer 1 prediction error loss 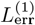, which measures the reconstruction error of input images. Minimization of the Layer 2 prediction error 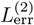 begins at epoch 15, and its weight in the objective function increases linearly through the end of training. This second error term corresponds to temporal predictions of input sequences—that is, expectations about the next image given previous presentations.

**Figure 3:**
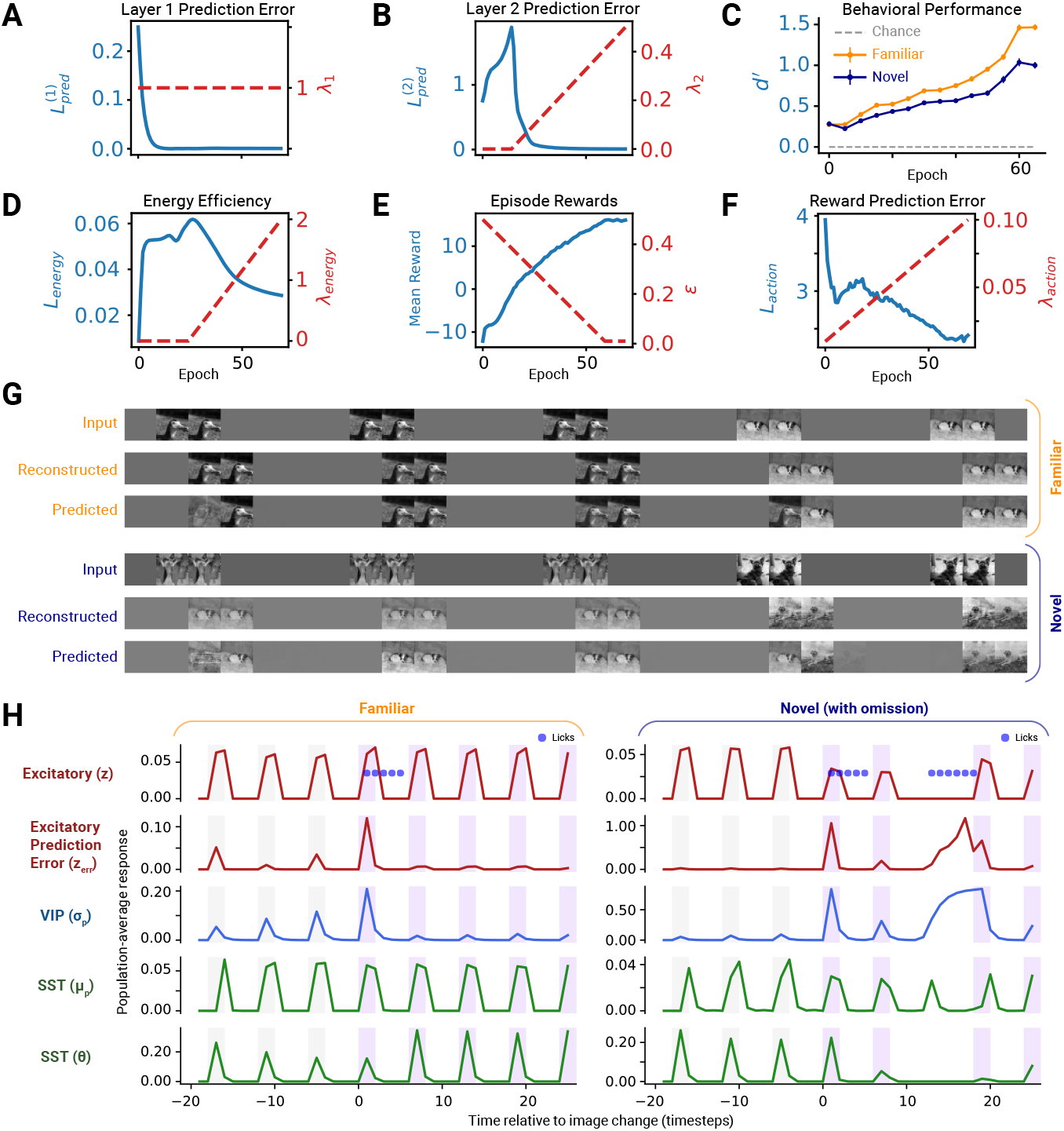
Training progress and example sequence responses. (A–F) Trajectories of different loss and behavioral metrics throughout training (solid blue traces showing average over 32 random seeds). Annealing schedules of the loss weights (*λ*_1_, *λ*_2_, *λ*_energy_, *λ*_action_) and the exploration probability (*ϵ*) are plotted with dashed red lines. To measure behavioral performance throughout training, we calculated *d*^*′*^ every 5 training epochs on both familiar and novel sequences with the current value of *ϵ*. We then averaged the *d*^*′*^ trajectories over 32 random seeds and plotted mean ± SEM. (G) Example familiar and novel input image sequences and the corresponding reconstructions and predictions from the model. Reconstructions are computed from the activities of Layer 2 representation (*z*) neurons after observing the input image with a one-time-step delay. Predictions are computed from higher area activities before an input image is observed and represent *expected* image presentations given sequence history. (H) Population-averaged responses for the different interneuron populations in Layer 2 (only experimentally recorded populations are shown) during an example familiar (left) and novel (right) sequence. Gray and purple bars represent presentations of different images. The novel sequence includes an omission in addition to an image change (gray to purple bars).

At the start of training, the energy efficiency loss *L*_energy_ is small but steadily increases until epoch 25, when the model begins explicitly optimizing it. This rise suggests the model is activating more *z* neurons to improve input reconstructions. At epoch 25, however, the model had already achieved asymptotically low reconstruction errors, so, by optimizing energy efficiency at this point, it prioritizes eliminating redundancy in its representations by making the activities of *z* neurons more sparse.

The model begins minimizing the reward prediction error *L*_action_ from the start of training, with a linearly increasing weight *λ*_action_. Reducing this error is crucial for learning accurate action-value estimates at each time step, which affects behavioral performance and total rewards. However, correctly learning these estimates requires collecting sufficient data about how different actions yield rewards in various contexts. Therefore, the model must explore more often early in training, when data is sparse. This is achieved by linearly annealing the exploration probability *ϵ* from 0.5 to 0.01 over the course of training. As *ϵ* decreases, the model collects more rewards by exploring less and reducing licks outside the change response window.

At each training epoch, we evaluate the model’s behavioral performance on both familiar and novel sequences by computing the signal-detection-theoretic measure *d*^*′*^. This evaluation is conducted using the current exploration probability *ϵ*, as determined by the annealing schedule. We categorize each image presentation as either a GO trial (change image) or CATCH trial (repeat image), and record whether the model licks within a 6-time-step window following the presentation. We compute the hit rate as the fraction of GO trials with a lick response, and the false alarm rate as the fraction of CATCH trials with a response. To avoid extreme values that produce infinite z-scores, we replace hit (false alarm) rates of 0 and 1 with values 1*/*2*N*_go_ (1*/*2*N*_catch_) and 1 − 1*/*2*N*_go_ (1 − 1*/*2*N*_catch_), respectively, where *N*_go_ (*N*_catch_) is the total number of GO (CATCH) trials ^56^. Finally, we compute

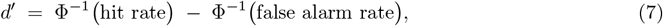

where Φ^−1^ is the inverse of the standard normal cumulative distribution function. The model shows significantly better performance than chance in both familiar and novel settings (Figure 3C), indicating it learns the behavioral task successfully and generalizes to novel sequences, albeit with slightly decreased performance.

To investigate the model’s learned representations of the task, we visualize how its internal predictions evolve over time in response to example image sequences (Figure 3G). At each time step *t*, the excitatory *z* neurons in the representation layer compute a reconstruction 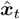 of the input image presented at the previous time step ***x***_*t*−1_. This is due to a single-time-step delay in the connections from the input layer to the intermediate and representation layers. At the same time step, the higher areas compute a prediction 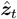 of the *z* neurons’ responses ***z***_*t*_. Notably,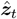 is computed from the higher area activities at time step *t* − 1, i.e. before the input ***x***_*t*−1_ is seen. Hence, it reflects the model’s expectation for ***z***_*t*_ given only previous information in the sequence. To compare these predictions with actual input images and their reconstructions, we project 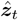 back into the input image space via the same feedback connections used to generate 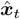, obtaining predicted images 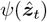. As shown in Figure 3G, the model accurately reconstructs familiar images but struggles with novel images, for which it lacks suitable representations. It also captures the temporal structure of the sequence: initially, it predicts a mixture of familiar images because it does not yet know which image will appear. After observing the first image, the model correctly predicts the sequence of familiar images until a new image is presented. When that change occurs, it briefly continues predicting the old image, then updates its prediction once the new image is seen.

Next, we examine the population-averaged activities of interneuron subpopulations in response to an example image sequence (Figure 3H). Both VIP and excitatory prediction error neurons show higher responses to novel than to familiar images, indicating that top-down predictions are less accurate and more uncertain for novel stimuli. Both populations also exhibit increased activity following image changes, signaling expectation violations. In contrast, excitatory *z* neurons and both SST subpopulations do not show a stereotyped response to image changes. While the excitatory *z* response appears to decrease after an image change in the novel sequence example (Figure 3H, right), this is due to their stronger image selectivity, as they primarily receive stimulus-tuned feedforward input. Consequently, this effect varies across sequences and disappears when averaged over multiple examples.

During omissions (Figure 3H, right), VIP and excitatory prediction error responses continue ramping up until the next image is presented. This effect was also observed in biological neurons ^23^, and has been hypothesized to arise from a predictive processing mechanism. Consistent with its role in encoding top-down predictions, the SST (*µ*^*p*^) population also exhibits a transient response after omission, reflecting an expected image presentation.

### Change and omission responses: model versus experiment

The primary purpose of our model is to show that a biologically plausible architecture that implements energy efficient predictive coding with reinforcement learning can produce three types of novelty effects in cortical interneurons: absolute, contextual, and omission. Rather than capturing precise biophysical dynamics, the model is designed to qualitatively replicate response patterns across interneuron populations by integrating normative computational objectives with a mechanistic circuit structure. Accordingly, we compare our model’s change and omission responses with experimental recordings from excitatory, VIP, and SST populations previously reported in Garrett et al. ^23,39^. These comparisons are primarily qualitative, as the model is not fit to experimental data and operates at an abstract level that differs in response scale. Nonetheless, they reveal notable similarities in the functional signatures of novelty processing (Figures 4 and 5).

**Figure 4:**
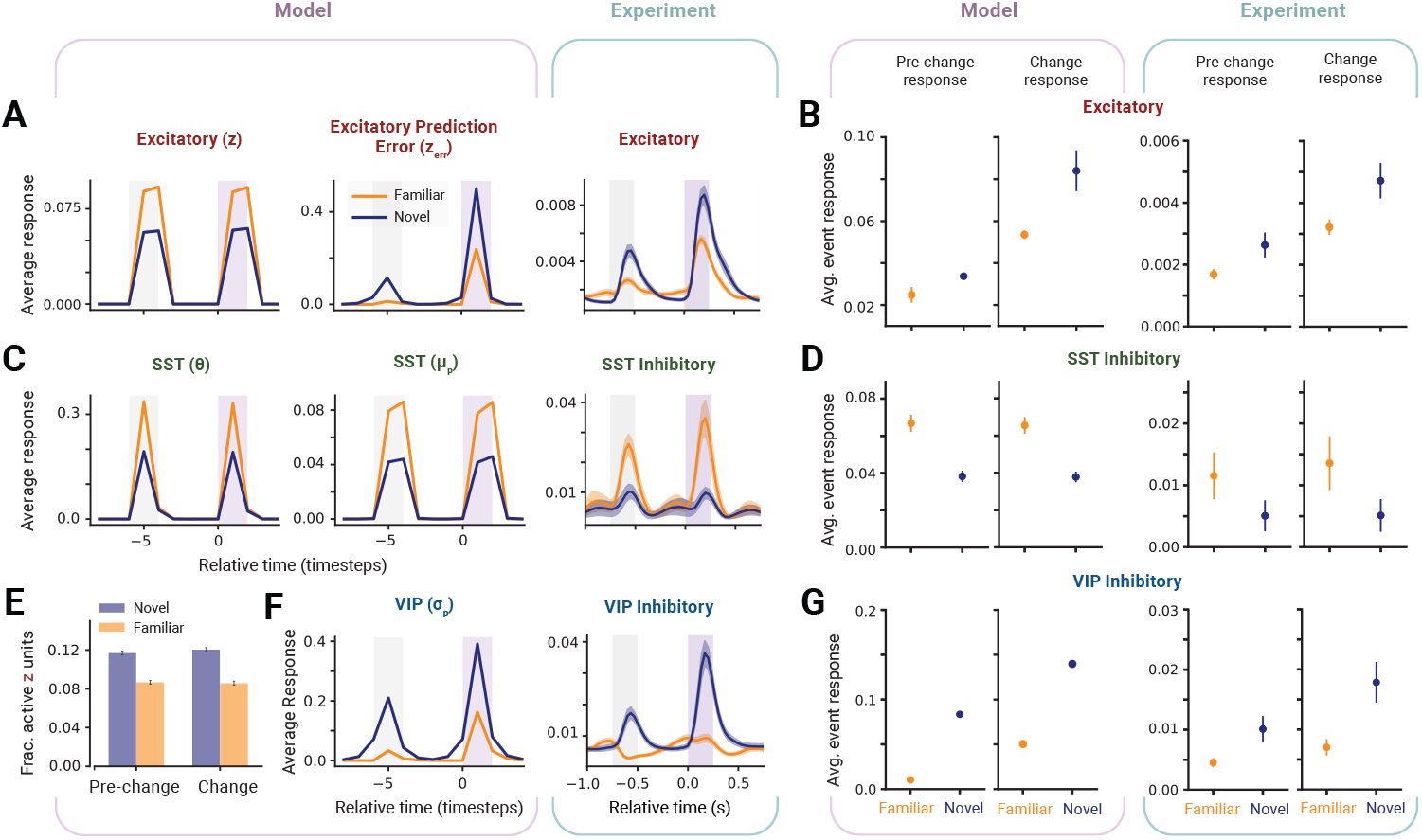
Model reproduces absolute and contextual novelty effects. (A) Responses of excitatory subpopulations in the model (left) and excitatory neurons in experiment (right, n=34 mice, 10,759 neurons) during image change trials. For both model and experiment, responses shown are averages across neurons and trials. For the model, those responses were further averaged across 32 training instances (random seeds). See Table S2 for the different population sizes and Methods for details on sequence (or trial) generation. For the experimental data, the number of trials varies across animals and sessions and can be obtained from the Allen Institute’s Visual Behavior (2P) dataset (link below). Shaded error bars in both cases represent SEM. (B) Average event-triggered excitatory responses for the model (left) and experiment (right). An event-triggered response for each neuron was computed as the mean response over a time window following the event (e.g. pre-change or change image presentation). Those responses were then averaged, for each neuron, across trials of different novelty conditions (familiar and novel), and used to compute the population mean ± 95% confidence intervals (CI). Average event-triggered responses from the two excitatory subpopulations in the model were concatenated to form a single distribution for the entire excitatory population. (C) and (D) same as (A) and (B) for the SST populations (n=15 mice, 417 neurons for the experimental data). (E) Average fraction of active neurons in the excitatory *z* subpopulations across trials and training instances. For each trial, average fraction was computed over a time window following the pre-change and the change image presentations. Error bars represent 99% CI. (F) and (G) Same as (A) and (B) for the VIP population (n=17 mice, 997 neurons for the experimental data). All experimental responses were reproduced using pre-processed data from the Allen Institute’s Visual Behavior (2P) dataset ^60^, found at https://doi.org/10.6084/m9.figshare.29207546.v2.

**Figure 5:**
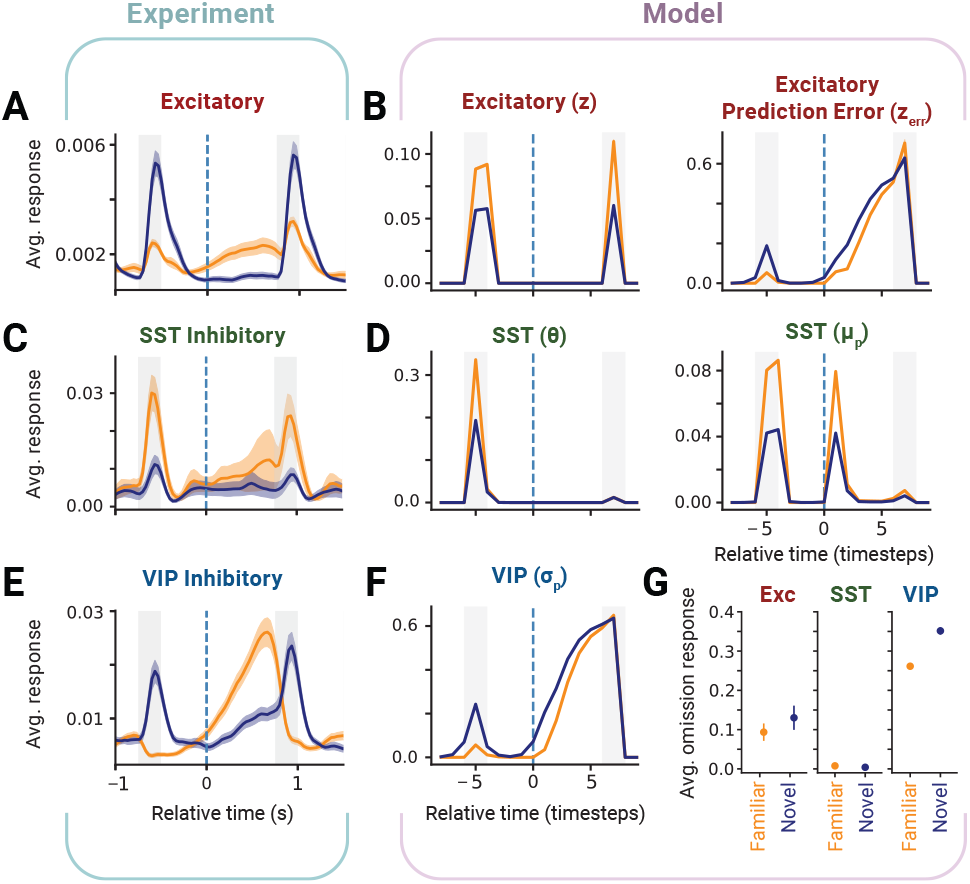
Model reproduces omission novelty effects. (A) Experimental response of excitatory neurons during omission trials. Dashed blue line indicates the time step when the expected image was omitted. Responses shown are averages across neurons and trials. Shaded error bars represent SEM. For the model, see Table S2 for population sizes and Methods for details on how omission trials are constructed. For the experimental data, details on the numbers of animals, cells, and trials are the same as in Figure 4. All experimental responses were reproduced using pre-processed data from the Allen Institute’s Visual Behavior (2P) dataset ^60^. (B) Model responses of the two excitatory subpopulations during omission trials. Responses shown are averages across neurons, trials, and training instances. Shaded error bars represent SEM. (C) Same as (A) for the experimental SST responses. (D) Same as (B) for the model SST subpopulations. (E) Same as (A) for the experimental VIP responses. (F) Same as (B) for the model VIP responses. (G) Average omission-triggered responses for the excitatory, SST, and VIP populations in the model. An omission-triggered response for each neuron was computed as the mean response over a time window following omission onset. Those responses were then averaged, for each neuron, across trials of different novelty conditions (familiar and novel), and used to compute the population mean ± 95% confidence intervals. To form a single distribution for each of the excitatory and SST populations, omission-triggered responses from their respective subpopulations were concatenated.

To visualize model responses during change and omission trials (Figures 4A,C,F and 5A-F), we took the mean of population-averaged responses across trials in all 32 instances (or random seeds) of training. For each image change sequence, the change trial started from 2 timesteps before the pre-change (or repeat) image to 2 timesteps after the change image. If the sequence contained an omission, the omission trial was taken from 2 timesteps before the pre-omission image to 2 timesteps after the post-omission image.

To quantify model responses in different novelty conditions (Figures 4B,D,G and 5G), we computed, for each neuron, an event-triggered response corresponding to the mean of that neuron’s response over a defined time window following an event (image presentation or omission). The window was four timesteps for images and six for omissions. We then calculated, for each neuron, the average event-triggered response across all trials of different event (pre-change, change, and omission) and novelty (familiar and novel) conditions. For each condition, we then created a distribution of average event-triggered responses across neurons within each cell population, from which we computed the mean ± 95% confidence intervals in Figures 4B,D,G and 5G. This analysis mirrors the approach used by Garrett et al. ^23^ to generate the corresponding experimental figures. Note that, since our model includes two excitatory and two SST subpopulations, we concatenated the average event-triggered responses from both excitatory subpopulations to form a single distribution for the excitatory population, and did the same for the SST population. While the subpopulations exhibit differences in response amplitude, such heterogeneity aligns with documented biological variability in both excitatory neurons and SST interneurons, which comprise diverse subtypes with distinct functional and electrophysiological properties ^23,57–59^. In Figure 4, we focus on the responses of populations for which experimental data are available; the responses of other areas in the model are shown in Figure S1.

### Absolute novelty

Within the excitatory population, prediction error neurons (*z*^err^) responded more strongly to novel images compared to familiar ones, whereas representation neurons (*z*) showed the opposite pattern (Figure 4A). The dynamics of the image-evoked responses are also different between the two subpopulations: prediction error neurons displayed a transient response, while representation neurons maintained a sustained response throughout the image presentation. When considering the event-triggered responses of the entire excitatory population, the overall activity was higher for novel images, aligning with experimental observations (Figure 4B). Additionally, the activity of *z* neurons was sparser for familiar images (Figure 4E), consistent with the model’s objective of maximizing energy efficiency.

Both SST subpopulations showed a stronger response to familiar compared to novel images, with the SST (*µ*^*p*^) neurons exhibiting a sustained image-evoked response and the SST (*θ*) neurons showing a more transient response (Figure 4C). Similar to experimental data, the SST population as a whole produced higher event-triggered responses for familiar than for novel images (Figure 4D). In contrast, VIP neurons responded more strongly to novel than to familiar images (Figure 4F), with their event-triggered responses exhibiting the same pattern (Figure 4G), in agreement with experimental findings.

### Contextual novelty

In response to image changes, the excitatory prediction error neurons show an increase in activity at the onset of change relative to pre-change responses (Figure 4A). This effect was not present in the representation neurons, which displayed the same image-evoked response to change and pre-change images. Consequently, the pooled event-triggered responses of the entire excitatory population were higher for the change image, agreeing with experiment (Figure 4B).

Similar to experimental data, there was no significant change response in either SST subpopulations (Figure 4C). The event-triggered response of the entire SST population was also similar for change and pre-change images (Figure 4D). On the other hand, VIP neurons responded more strongly to change images relative to pre-change in both novel and familiar trials (Figure 4F). This effect was also reflected in their average event-triggered responses (Figure 4G).

### Omission novelty

Garrett et al. ^23^ showed that VIP neurons exhibit a unique ramping effect during omissions of expected stimuli, which was present to a lesser extent in excitatory neurons and absent in SST neurons (Figure 5A,C,E). Additionally, this effect was strong when familiar images were omitted but was significantly reduced for novel images. In our model, VIP neurons also displayed a similar ramping response during omissions, which was suppressed by the subsequent image presentation (Figure 5F). This ramping response was present in the excitatory prediction error subpopulation but absent in the excitatory representation neurons and the SST (*θ*) subpopulation (Figure 5B,D). SST (*µ*^*p*^) neurons responded transiently after image omission (Figure 5D), indicating that the model expected an image presentation at that time step. To quantify these effects, we show the average event-triggered responses for the entire excitatory and SST populations (pooled across subpopulations) and the VIP population. Similar to experimental observations, the omission ramping response was strongest in VIP neurons, weaker in excitatory neurons, and absent in SST neurons (Figure 5G). However, in contrast with experiment, omission responses were stronger for novel images in VIP neurons but were not significantly different between novel and familiar images in excitatory neurons.

### Role of reinforcement learning

To investigate the role of reinforcement learning in shaping population responses, we test a *perception-only* version of our model that does not have the components for reward prediction and behavioral learning. More specifically, this version does not include the Reward area in the higher-level processing layer, and does not implement action selection or reward-based feedback modulation. Everything else is kept the same as in the full model, including training, evaluation, and hyperparameter settings.

In the perception-only model, absolute novelty effects in all populations were similar to those in the full (perception-action) model (Figure S2). However, contextual novelty effects, or change responses, disappeared in the VIP population and were reduced in the excitatory prediction error subpopulation (Figure 6A). For VIP neurons, in both familiar and novel conditions, population-averaged responses were the same for pre-change and change images as were the distributions of average event-triggered responses across neurons (Figure 6A, bottom). For excitatory prediction error neurons, change responses were still present but were much weaker during novel sequences (Figure 6A, top).

**Figure 6:**
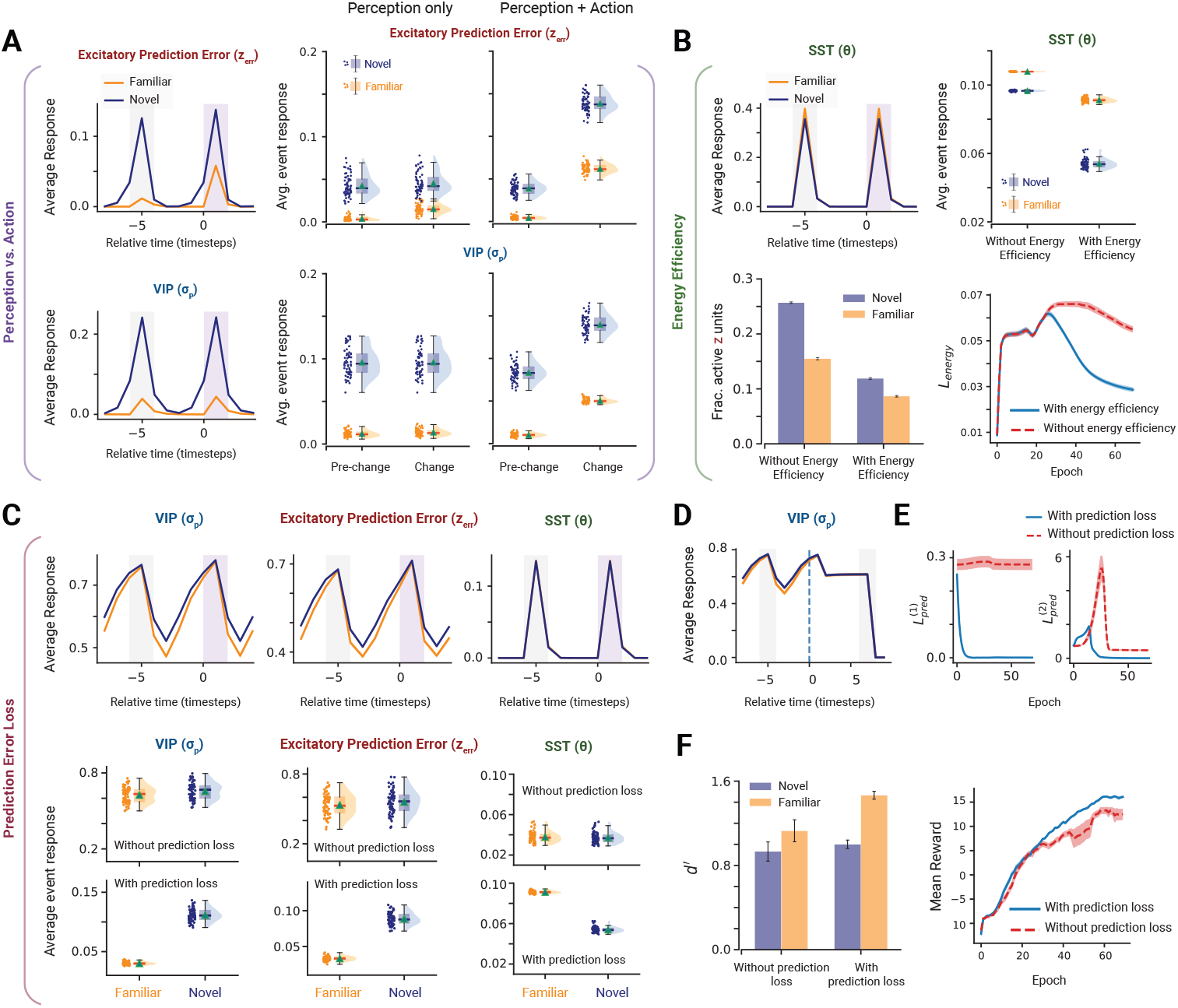
Effects of different computational objectives on model responses. (A) Image change responses of the excitatory prediction error (*z*^err^) subpopulation (top left) and the VIP population (bottom left) *without* the reinforcement learning component (perception only). On the right, average event-triggered responses are shown for each population as raincloud plots ^61^ in the perception-only case as well as the full model (perception + action). Each dot represents the event-triggered response of a neuron in the respective population averaged across trials of different event types (pre-change and change) and experience levels (familiar and novel). Removing the reinforcement learning component abolishes contextual novelty effects in VIP neurons and diminishes them in the excitatory prediction error neurons. (B) Impact of removing the energy efficiency objective. Top-left: image change responses of SST (*θ*) neurons. Top-right: raincloud plots of SST (*θ*) event-triggered responses, averaged across pre-change and change image for each of the familiar and novel conditions. Removing the energy efficiency objective diminished the absolute novelty effects in SST (*θ*) neurons. Bottom-left: fraction of active neurons in excitatory *z* subpopulation, averaged across trials and (pre-change and change) image presentations, with and without the energy efficiency objective. Error bars represent 99% CI. Bottom-right: training trajectories of the energy efficiency loss function, average across training instances. Shaded error bars represent SEM. (C-F) Impact of removing the predictive coding objectives. (C) Top: Image change responses of VIP, excitatory prediction error (*z*^err^), and SST (*θ*) neurons without the prediction error loss. Middle and Bottom: raincloud plots of event-triggered responses for each population with (middle row) and without (bottom row) the prediction error losses in both Layer 1 and Layer 2. Responses are averaged across pre-change and change images for each of the familiar and novel conditions. Removing the predictive coding objective abolishes absolute novelty effects. (D) VIP Omission response without the prediction error loss. (E) Training trajectories of Layer 1 and Layer 2 prediction error losses. Note the sharp decrease in the Layer 2 prediction error loss around epoch 25, when the energy efficiency optimization begins, leading to sparser excitatory activity and thus lower prediction error at Layer 2. (F) Left: d-prime values with and without the prediction error loss, averaged across model instances (n = 32 random seeds). Error bars represent 95% CI. Right: average reward accumulated per sequence throughout training.

These results suggest that reinforcement learning and reward modulation are necessary to produce the contextual novelty effects in VIP neurons. For excitatory neurons, sensory expectation violations alone can still give rise to change responses, but reinforcement learning significantly enhances those effects.

### Role of prediction error loss

A key principle of the predictive coding framework, and its biological implementations, is the minimization of prediction errors at different layers in the cortical hierarchy. To investigate the impact of prediction error minimization on population responses, we trained the model without prediction error losses at Layers 1 and 2 by setting *λ*_1_ = *λ*_2_ = 0 throughout training. As expected, Layer 1 prediction error 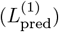 did not decrease throughout training compared to the full model (Figure 6E, left). In contrast, Layer 2 prediction error 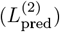 continued to increase until epoch 25, when the model began optimizing the energy efficiency loss (Figure 6E, right). At this point, as *z* neuron activity became sparser, 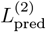 decreased sharply but plateaued at a higher value than in the full model for the remainder of training. Responses of populations affected by the removal of the prediction error objective are shown in Figure 6C–F; responses of other populations in the model are shown in Figure S4.

Removing the prediction error loss impacted absolute novelty effects in excitatory prediction error (*z*^err^), VIP, and SST (*θ*) neurons (Figure 6C), eliminating significant differences between responses to familiar and novel images. Moreover, VIP and prediction error neurons no longer exhibited enhanced responses to image changes, nor were their response dynamics time-locked to image presentations. This likely reflects their reliance on top-down feedback from the higher areas, which, in the absence of prediction error minimization, failed to encode the temporal structure of the task. Additionally, the omission response of VIP neurons was disrupted (Figure 6D), as they no longer exhibited the ramping activity observed during omissions in the full model.

Eliminating the prediction error loss also impaired task performance, particularly on familiar trials. Change detection performance, measured by *d*^*′*^, was significantly worse for familiar trials compared to the full model (Figure 6F, left). Furthermore, the model accumulated fewer rewards over training when prediction error was not minimized (Figure 6F, right). These findings suggest a link between building predictive representations and achieving good task performance.

### Role of energy efficiency

To examine the impact of energy efficiency on population responses, we trained the model without the energy efficiency objective, ∥ ***z*** ∥_1_, by setting *λ*_energy_ = 0. The response patterns of excitatory, VIP, and SST neurons remained largely unchanged from Figure 4 and 5, with absolute, contextual, and omission novelty effects persisting despite the removal of the energy efficiency constraint (see Figure S3). However, the absolute novelty effects in SST (*θ*) neurons were significantly diminished, as the difference in their responses to familiar and novel images was markedly reduced compared to the energy-efficient model (Figure 6B, top). This indicates that the model utilizes the SST (*θ*) population when optimizing energy to achieve more efficient representations of familiar images.

Additionally, removing the energy efficiency constraint led to a substantial decrease in the sparsity of excitatory *z* neuron responses under both novel and familiar conditions, reflecting greater redundancy and higher energetic costs (Figure 6B, bottom left). Interestingly, while explicitly optimizing energy efficiency led to a more pronounced reduction in its loss value, a slight decrease was still observed even when the objective was not enforced (Figure 6B, bottom right). This suggests that the underlying circuitry itself, particularly the VIP-SST-excitatory pathway, contributes to optimizing energy efficiency even in the absence of an explicit computational objective. We explore this effect further in the following section.

### Interplay between disinhibition and energy efficiency

To investigate the connection between disinhibition and energy efficiency, we examined the effects of ablating different connections within the VIP-SST-Excitatory circuit.

First, we ablated the *θ*-*z* connection, effectively removing inhibition from the SST (*θ*) population onto the excitatory (*z*) neurons (Figure 7A). This ablation produced effects on SST (*θ*) responses that closely resembled those observed when removing the energy efficiency objective, despite energy efficiency still being explicitly optimized. Specifically, the difference between responses to familiar and novel images was diminished (Figure 7B). Additionally, excitatory prediction error neurons exhibited an overall increase in activity after removing the *θ*-*z* connection (Figure 7C). Notably, a similar but smaller increase in prediction error responses was observed when the energy efficiency loss was removed while this connection remained intact. These findings suggest that the disinhibitory circuit inherently contributes to energy efficiency optimization, even in the absence of an explicit computational objective. Moreover, when the *θ*-*z* connection was ablated, the model was unable to minimize the energy efficiency loss, as reflected in the loss trajectories throughout training (Figure 7D). This highlights the necessity of this inhibitory pathway for optimizing representational efficiency. The novelty effects in other population responses under this ablation remained largely the same (Figure S5).

**Figure 7:**
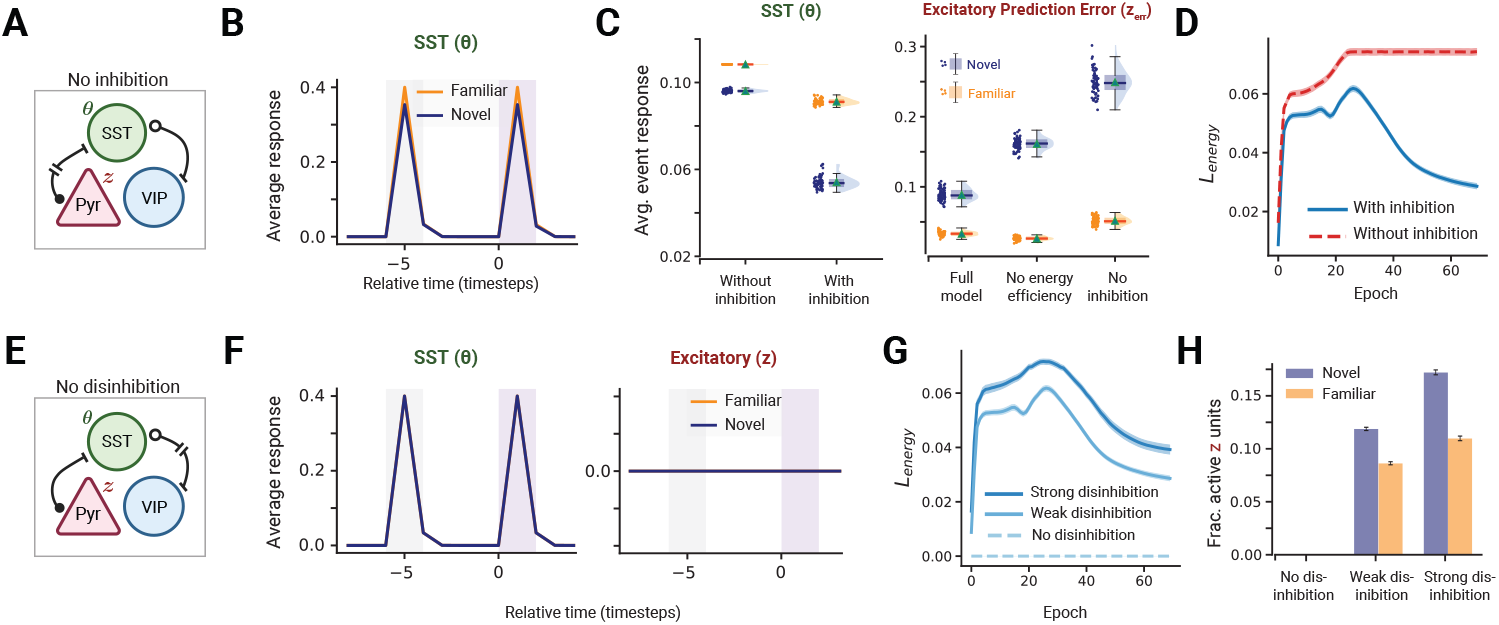
VIP-SST disinhibitory circuit balances energy efficiency and representational capacity. (A) Schematic of the VIP-*θ*-*z* circuit in our model, with the *θ*-*z* connection ablated, corresponding to no inhibition of *z* neurons. (B-D) Impact of no inhibition. (B) Image change responses of SST (*θ*) neurons. (C) Raincloud plots of SST (*θ*) and prediction error (*z*^err^) event-triggered responses, averaged across pre-change and change images for each of the familiar and novel conditions. Without the *θ*-*z* connection, absolute novelty effects in SST (*θ*) neurons are diminished and prediction error activities (*z*^err^) increase. (D) Training trajectories of the energy efficiency loss function with and without the *θ*-*z* connection, averaged across training instances. Shaded error bars represent SEM. (E) Schematic of the VIP-*θ*-*z* circuit in our model, with the VIP-*θ* connection ablated, corresponding to no disinhibition of *z* neurons. (F-H) Impact of no disinhibition. (F) Image change responses of SST (*θ*) and excitatory representation (*z*) neurons. Without disinhibition, SST (*θ*) no longer distinguish between familiar and novel images and excitatory (*z*) neurons are silenced due to significantly low E/I ratio. (G) Training trajectories of the energy efficiency loss function with varying VIP-*θ* connection strengths. (H) Fraction of active *z* neurons, averaged across trials and (pre-change and change) image presentations, with different VIP-*θ* connection strengths. Error bars represent 99% CI.

Next, we examined the effects of ablating the VIP-SST connection (Figure 7E) by systematically varying its strength 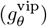: 0 (no disinhibition), 2 (weak disinhibition), and 4 (strong disinhibition). When the connection was completely removed (no disinhibition), excitation-inhibition (E/I) balance was severely disrupted, leading to the complete silencing of excitatory (*z*) neuron activity (Figure 7F) and impairments in all other population responses (Figure S6), ultimately degrading behavioral performance. Additionally, absolute novelty effects in the SST (*θ*) population were abolished, with responses becoming solely driven by untuned feedforward input. These results imply a possible role for disinhibition in regulating E/I balance, which is essential for meaningful cortical computations.

Furthermore, varying VIP-SST disinhibition strength revealed a role in regulating energy efficiency optimization. As disinhibition increased, the energy efficiency loss objective also increased, indicating reduced sparsity in excitatory (*z*) neuron activity due to weaker SST inhibition (Figure 7G,H). This suggests that disinhibition may be involved in modulating the trade-off between representational power and energy efficiency, balancing energetic cost with the ability to encode information.

### Comparison to a response adaptation model with Hebbian learning

Could the observed novelty effects be accounted for by simple biological mechanisms such as response adaptation and Hebbian learning? To address this question, we constructed a mechanistic firing-rate model of the cortical microcircuit that incorporates both response adaptation and Hebbian plasticity (see Methods). The model includes excitatory, VIP, and SST populations, with connectivity mirroring that of our original model (Figure 8C). We evaluated the model under the same task conditions as in our main simulations and assessed novelty effects both with and without Hebbian learning (Figure 8). Population-averaged response traces and event-triggered responses were computed using the same procedures as in the original model.

**Figure 8:**
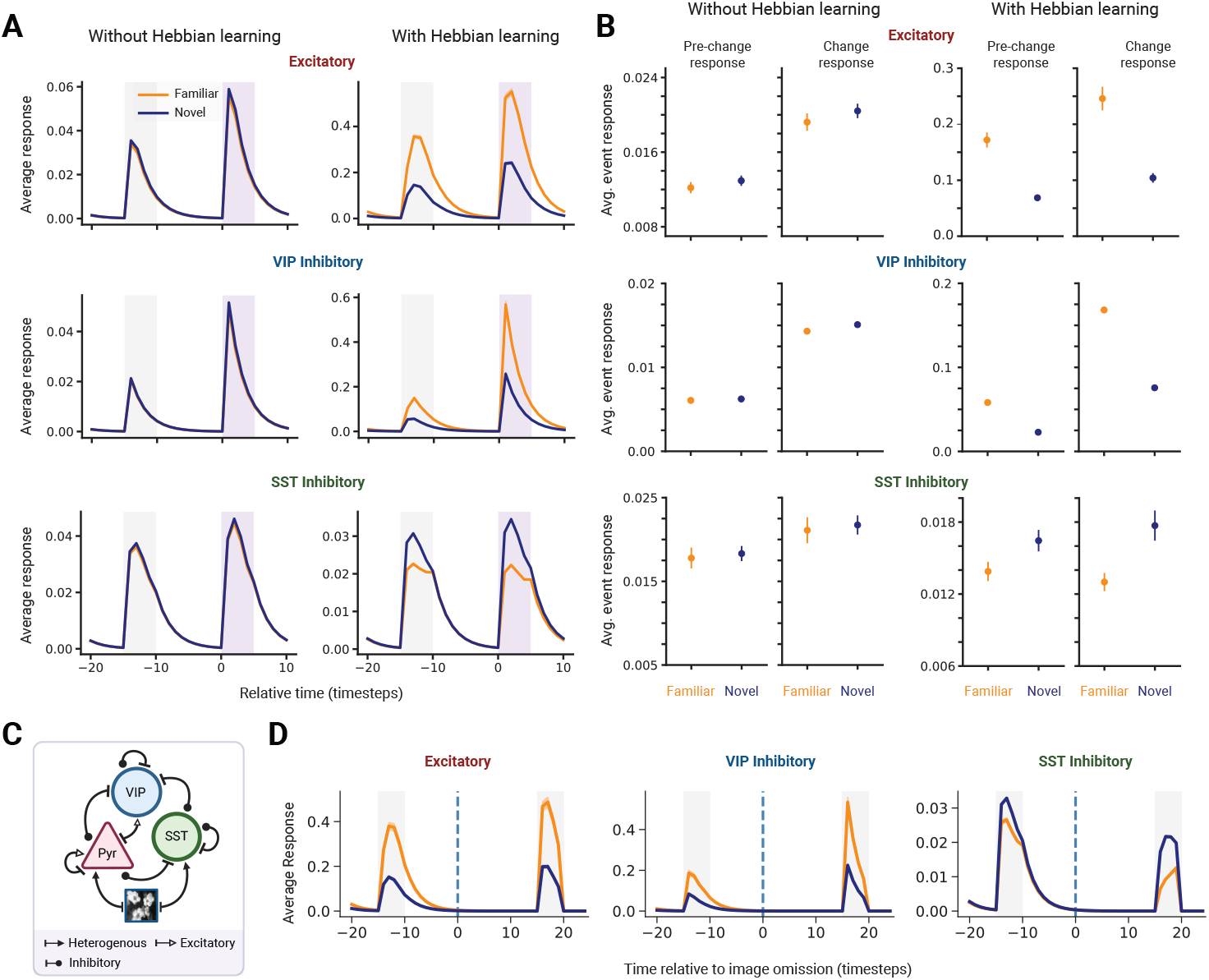
Novelty effects in a model with response adaptation and Hebbian learning. (A) Image change response of excitatory, VIP, and SST neurons in the adaptation model with (right) and without (left) Hebbian learning. (B) Average event-triggered responses for each population with and without Hebbian learning across different event types (change vs. pre-change) and novelty conditions (familiar vs. novel). Event triggered responses were computed using the same procedure as in the original model. Response adaptation alone can capture contextual novelty effects, while Hebbian learning produces absolute novelty effects that are opposite to experimental observations. (C) Schematic showing the connectivity of the interneuron circuit in the adaptation model. (D) Omission responses of excitatory, VIP, and SST neurons in the adaptation model with Hebbian learning.

In the absence of Hebbian learning, the adaptation model exhibited no differences in population responses to familiar versus novel images (Figure 8A,B), as expected given the lack of a training phase to bias responses toward a specific image set. However, by tuning the adaptation strengths of different populations, the model could reproduce contextual novelty effects observed experimentally. Specifically, assigning higher adaptation strengths to excitatory and VIP neurons led to increased responses to image changes relative to pre-change responses. In contrast, SST neurons—with weaker adaptation—did not display significantly stronger responses to image changes.

We next introduced Hebbian learning by training the model on familiar image sequences for 20 epochs, during which synaptic weights were updated using a Hebbian rule (see Methods). After training, we examined the model’s responses during familiar and novel change trials (Figure 8A,B). Hebbian plasticity produced absolute novelty effects across all three populations, but in the direction opposite to that observed experimentally: excitatory and VIP neurons responded more strongly to familiar than to novel images, while SST neurons—subject to strong inhibition from VIP neurons—responded more strongly to novel images. These effects are consistent with Hebbian mechanisms, in which synaptic strengthening occurs for circuits repeatedly activated during familiar image presentations, resulting in elevated responses to those inputs.

Finally, the adaptation model failed to produce omission responses, even when Hebbian learning was included (Figure 8D). This contrasts with experimental findings and our original model, both of which exhibit ramping activity in VIP and excitatory neurons during stimulus omission.

## 3 Discussion

We present a biologically plausible cortical circuit model that integrates predictive coding, energy efficiency, and reinforcement learning objectives to account for diverse novelty-related responses in the visual cortex. By assigning computational roles within this framework to different interneuron populations, our model reproduces three types of novelty effects observed experimentally in VIP, SST, and excitatory neurons ^23^. We show that the predictive coding objective is necessary for the emergence of absolute novelty effects (novel versus familiar) in all three interneuron populations. Our model also captures omission novelty in VIP neurons (ramping during stimulus omission), but fails to do so without the predictive coding objective. Incorporating reinforcement learning and reward-based modulation leads to contextual novelty effects (change versus pre-change) in VIP neurons and amplifies the change responses of excitatory neurons. Finally, we demonstrate an important relationship between the VIP-SST disinhibitory pathway and energy efficiency, suggesting that subpopulations of SST neurons may be involved in implementing efficient coding schemes at the circuit level ^1^.

Besides replicating experimental responses, our model provides a circuit-level theory for how predictive coding may be implemented in cortical microcircuits with different interneuron subtypes, extending existing theories which abstract away inhibitory interneuron diversity ^4,5,62^. In this theory, each interneuron population represents an algorithmic node of the predictive coding framework. At least two functional classes of neurons are required to implement predictive coding: representation neurons and prediction-error neurons ^3,63^. These correspond to the two subpopulations of excitatory neurons in our model, *z* and *z*^err^, respectively. As such, representation neurons (*z*) encode latent states (or causes) of sensory observations, which are computed during inference by integrating stimulus-driven, feedforward input with local inhibitory input from PV and SST (*θ*) neurons. Feedforward input computes the mean of the posterior distribution over latent states given observations. This design is consistent with the strong tuning properties of excitatory neurons since feedforward input tends to be highly tuned for specific stimulus features. PV neurons also receive feedforward input and represent the precision of the posterior distribution, in line with evidence for their role in feedforward inhibition and gain control of sensory inputs ^11–13,64–66^. By combining divisive PV inhibition with feedforward input, representation neurons in our model effectively *sample* from the posterior distribution. This supports the neural sampling hypothesis ^67,68^, which posits that neural firing rates represent samples from probability distributions such as the Bayesian posterior. In our implementation, local excitatory-inhibitory interactions perform this sampling operation to generate probabilistic representations. A similar mechanism also enables sampling from the prior, by integrating SST (*µ*^*p*^) and VIP input to the prediction error neurons.

Prior expectations, corresponding to top-down predictions from higher areas^3,5,69^, are encoded by two inhibitory interneuron populations: VIP neurons encode the prior distribution’s variance, thus representing the uncertainty of predictions, and SST (*µ*^*p*^) neurons encode the prior distribution’s mean. These roles are consistent with experimental evidence that VIP and SST neurons receive mainly top-down feedback signals ^11,18,19^, but they also make sense computationally since top-predictions must have a net inhibitory effect to allow the calculation of prediction errors at lower layers ^4,5,63,69^. Moreover, evidence from experimental and modeling studies suggests a role for VIP neurons in contextual modulation and uncertainty representation. For example, in the mouse primary visual cortex, VIP neurons are activated by weak (low-contrast) stimuli and during locomotion, potentially signaling situations of higher sensory uncertainty ^70^. This could imply that VIP activity increases when prior predictions are less certain, adjusting the network to enhance sensitivity. Additionally, neuromodulatory and feedback signals routed through disinhibitory pathways (such as to VIP) may be encoding uncertainty (or precision) associated with top-down predictions ^64,69,71^. Computational evidence has also suggested that VIP neurons contribute to mismatch responses by amplifying differences between feedforward and feedback inputs ^7,72^, consistent with their role in modulating prediction errors in our model. The role of SST neurons in encoding prior expectations is also supported by evidence showing they gate top-down input through subtractive inhibition of excitatory neurons ^65,66,73^. SST neurons have also been causally linked to surround suppression ^18,74,75^ which can also be explained in a predictive coding framework ^3,6^.

Recent models have taken important steps toward integrating interneuron diversity into predictive coding frameworks ^8,76,77^. Hertag et al. ^8^ demonstrate how distinct prediction-error neurons can emerge through inhibitory plasticity in circuits with multiple interneuron types. While this model emphasizes the role of E/I balance, it relies on abstract prediction signals in a fully predictable sensory environment, lacking an explicit computational objective. Our model complements this by assigning normative roles to cell-type-specific subpopulations, grounded in biological connectivity and task-driven learning. Lee et al. ^76^ present a cortical circuit model of predictive coding with multiple cell types, realistic connectivity, and emergent mismatch responses; however, they do not explore how distinct interneuron subtypes map onto algorithmic components. Wilmes et al. ^77^ implement a Bayesian approach in which layer 2/3 pyramidal neurons compute uncertainty-modulated prediction errors, with SST neurons encoding the mean of the prior distribution and PV neurons encoding input variance. However, their model does not include VIP neurons nor does it account for the uncertainty associated with top-down predictions. These models emphasize either circuit mechanisms or error minimization, using simplified, non-behavioral settings with artificial inputs, which limits their comparability to experimental data. In contrast, our approach integrates a multi-objective normative framework with a biologically grounded circuit model that encompasses all major cortical cell types, offering a unified account of how predictive coding, energy efficiency, and reinforcement learning map onto distinct cell-type-specific mechanisms. Moreover, by training the model on a naturalistic, task-driven paradigm, we enable more direct and faithful comparisons to biological responses.

A key feature of the experimental data from Garrett et al. ^23^ is the diversity of novelty responses observed within each cell class, including excitatory, VIP, and SST populations. This diversity is reflected in our model through its explicitly defined subpopulation structure. Rather than treating each population as homogeneous, we assigned distinct computational roles to subgroups within both excitatory and SST neurons, enabling the model to replicate the response heterogeneity observed in those populations. For instance, the experimental work identifies excitatory clusters that respond more strongly to novel images, similar to prediction error neurons in our model, and others with higher responses to familiar images, similar to representation neurons. Therefore, our model offers a theoretical interpretation of this differentiation: excitatory neurons tasked with signaling prediction errors are more active during novel trials, while those dedicated to representing input features are preferentially engaged during familiar trials. Similarly, the SST population in the experiment contains subclusters that exhibit no response to stimulus omissions and others that show weak omission-triggered activity. This differentiation is recapitulated in our model which divides SST into two subpopulations with distinct functional roles: SST (*θ*) neurons, which facilitate efficient coding and do not respond to omissions, and SST (*µ*^*p*^) neurons, which carry top-down predictions signals and respond transiently when an expected stimulus is omitted. By explicitly embedding this subpopulation structure into the circuit, the model accounts for the within-type diversity of response profiles observed experimentally and highlights the computational need for such differentiation in implementing energy-efficient predictive coding.

Our model shows that contextual novelty effects in VIP neurons are a consequence of reinforcement and reward-based feedback modulation. In excitatory neurons, although these effects were amplified by reinforcement learning, predictive coding was sufficient for them to emerge, particularly in familiar settings. Biologically, this implies that VIP activation in contextually novel settings depends on top-down signals tied to behavioral relevance. Indeed, multiple experimental findings suggest that VIP neurons are activated by reinforcement signals ^13,17^ and expected rewards ^78,79^, which could enhance pyramidal cell responses in reward contexts. In a recent study analyzing the same dataset of Garrett et al. ^23^, Piet et al. ^40^ showed that mice using a visual strategy (attending to image changes for reward) exhibited greater VIP activity than mice relying on timing. Computational evidence also points to VIP neurons as mediators of reward-based top-down plasticity ^80^.

One novel insight from our model is the interplay between disinhibition and energy efficiency in regulating cortical responses. The VIP-SST pathway inherently implements a form of gain control that balances information encoding against metabolic cost. In our model, SST (*θ*) inhibit excitatory representation neurons (*z*), promoting sparsity, while VIP neurons can relieve this inhibition by inhibiting SST (*θ*) neurons. Interestingly, we found that even without an explicit energy efficiency objective, representation sparsity slightly increased throughout training, which no longer occurred when the connection between SST and excitatory neurons was ablated. This implies that the VIP-SST disinhibitory motif naturally acts as a circuit mechanism for sparse and efficient coding^1,2^. Our ablation analyses in Figure 7 also demonstrate a role for this motif in regulating E/I balance. Removing the VIP → SST (*θ*) connection silenced excitatory neurons, indicating that a baseline level of disinhibition is essential to prevent over-inhibition. Conversely, increasing disinhibition reduced representation sparsity. A unifying interpretation of these results is that the VIP-SST disinhibitory motif implements a circuit-level trade-off between representational power and sparse coding, which can be controlled by modulating synaptic strengths through local plasticity changes. This control can occur over long time scales through experience, resulting in the differentiation of responses to familiar and novel stimuli, and over shorter time scales according to task demands. Although experimental studies are needed to directly verify this interpretation, there is some current evidence supporting it. For example, Liang et al. ^81^ showed that the development of sparse representation in layer 2/3 was experience-dependent, accompanied by changes in the distribution of E/I ratios, and can be modulated by manipulating local inhibitory neurons. Sparse coding is also thought to underlie the phenomenon of surround suppression ^82,83^, which has been shown to involve SST neurons ^18,74,75^ and engage the VIP-SST disinhibitory pathway ^84^.

Our work is part of a growing effort to model experimentally observed, cell-type-specific novelty responses as shown in Garrett et al. ^23^. Most relevant to our setting is the recent model by Aitken et al. ^20^, which reproduces a wide range of novelty effects in VIP neurons using an unsupervised Hebbian-like learning approach. They introduced familiarity-modulated synapses (FMS) that weaken or strengthen based on a neuron’s history of activation. By incorporating synapses with multiple such plasticity mechanisms, their model could fit absolute, contextual, and omission novelty responses, even capturing heterogeneity within the VIP population. This is an elegant demonstration that relatively simple synaptic modulations can give rise to complex novelty responses, reinforcing the idea that the brain may not need explicit “novelty detectors” but can adapt circuits to produce these effects. However, the FMS model only reproduces novelty effects in VIP neurons. To capture these effects in other populations, different FMS mechanisms would need to be implemented for other synaptic connections in the network ^20^, increasing the model’s complexity as more effects are modeled. Furthermore, the FMS model, similar to our baseline adaptation model, is mechanistic; it shows that certain plasticity mechanisms can produce novelty responses but it does not explain why interneuron subtypes should behave as they do nor why biological connections should develop the specific FMS mechanisms used in the model. Our model, by contrast, starts from normative principles and shows that assigning algorithmic roles to interneuron subtypes and optimizing specific computational objectives naturally leads to the observed patterns. The mechanistic and normative approaches are complementary: the mechanistic models demonstrate feasibility with biologically plausible mechanisms, while our model offers a higher-level interpretation linking those mechanisms to computational goals.

Our model makes several experimentally testable predictions. First, because VIP change responses are absent without reinforcement learning, our model predicts that VIP neurons do not respond strongly to deviant stimuli in passive (unrewarded) oddball paradigms, but excitatory neurons do. While numerous findings support this prediction for excitatory neurons^33,85^, VIP neurons have not been thoroughly studied in oddball experiments ^86^. However, a recent study by Bastos et al. ^14^ employing a passive paradigm showed that VIP neurons indeed do not exhibit a deviance detection response, in line with our prediction. Second, although we have focused on the responses of excitatory, SST, and VIP neurons (from which experimental recordings were obtained), our model includes PV neurons and shows that they also exhibit novelty effects (Figure S1). Specifically, we predict that PV neurons respond more strongly to novel stimuli compared to familiar ones. This prediction is already supported by multiple experimental findings that PV neurons are activated by novel stimuli and suppressed with experience as those stimuli become familiar ^21,87,88^. Third, because the VIP-SST disinhibitory motif is a canonical circuit found throughout the cortex ^89^, our findings have implications that extend beyond *visual* change detection. We predict that VIP and SST neurons in non-visual cortical areas exhibit novelty effects similar to those captured in our model but for different sensory modalities. Finally, ablating different connections in the VIP-SST-excitatory circuit directly affected the familiar and novel differences in SST responses. Therefore, we predict that conducting similar manipulations in vivo, for example by optogenetically silencing VIP neurons, the absolute novelty effects in SST neurons will be significantly diminished. We do not predict that they will completely disappear since other subpopulations not inhibited by VIP neurons, such as SST (*µ*^*p*^) in our model, may still exhibit such effects.

### Limitations and future directions

Our model abstracts away some biophysical aspects of neural responses and anatomical details of real cortical circuits. For example, we do not explicitly model spiking and firing rate dynamics, which causes the shape and scale of our model responses to be less biophysically realistic. We chose to focus on *qualitatively* reproducing the effects observed experimentally rather than a direct quantitative comparison to experimental measurements, which would require modeling biophysical dynamics and fitting them to data. Additionally, we do not explicitly model the multilayer structure of cortical areas beyond a single local circuit level. Future work could involve replicating this local circuit across multiple layers in a laminar structure while incorporating biophysically realistic firing rate dynamics.

Our model does not capture the familiar versus novel differences in VIP omission responses, which were reported to be stronger for familiar than novel omissions ^23^. This could be a consequence of VIP neurons solely representing the uncertainty of top-down predictions, which leads to the disentanglement of omission and absolute novelty effects in their responses. Specifically, omissions increase uncertainty with respect to the temporal structure of the sequence while image novelty increases uncertainty with respect to the visual input itself. In our model’s VIP responses, the two effects are disentangled and combine to produce higher omission responses in the novel condition. However, in the brain, VIP activity is likely modulated by more complex interactions with local and feedback signals, making their uncertainty representation about temporal regularity dependent on image familiarity. This would also explain why VIP activity ramps up before familiar image presentations but not before novel ones, a phenomenon observed experimentally but not captured by our model ^23,39^.

## 4 Methods

### Notation

We use bold uppercase letters, e.g. ***W***, to refer to matrices, and bold lowercase letters to refer to vectors, e.g. ***z***. To refer to the weight matrix parameterizing connections from population *a* to population *b*, we use the notation ***W***_a→b_. Finally, we use the notation ***I*** to refer to the identity matrix.

### Change detection task

#### Experimental setup

The experimental change detection task in Garrett et al. ^23^ consists of a series of image presentations, where one image is repeated for a variable number of times until a change occurs and a different image is presented. Images were presented for 250 ms and interleaved with a gray screen 500 ms in duration (Figure 1E). Mice were trained to detect image changes by licking for a reward within a 750-ms response window following the change (Figure 1E). Experiments were conducted over two stages: behavior training then 2-photon imaging (Figure 1D). During behavioral training sessions, mice learned the task with a set of natural scene images (Figure 1A) that gradually became a *familiar* set over thousands of presentations. After reaching a sufficient performance threshold, mice performed the task during 2-photon imaging sessions, which consisted of sessions with familiar images, followed by multiple sessions with *novel* images (i.e, not encountered during behavioral training sessions). Garrett et al. ^23^ also ran additional “Novel+” sessions in which these initially novel images were presented after becoming familiar over time. Because our analysis focuses strictly on familiar and novel conditions, we exclude neural data from these Novel+ sessions when reproducing experimental responses in Figures 4 and 5. The task also included *omission* trials, where non-change images are unexpectedly omitted (i.e., replaced by a gray screen) with 5% probability (Figure 1E, bottom). See Garrett et al. ^23^ for a more detailed description of the task and training procedure.

Neural activity was recorded using 2-photon calcium imaging in the mouse visual cortex. Recordings were obtained from transgenic mice expressing the calcium indicator GCaMP6f in excitatory neurons, somatostatin (SST)-expressing inhibitory neurons, and vasoactive intestinal peptide (VIP)-expressing inhibitory neurons. Cells were imaged in both primary (VISp) and secondary (VISl) visual cortical areas. For more details on the experimental dataset and imaging procedure, see Garrett et al. ^23^.

#### Model setup

Our setup mirrors the experimental change detection task described above. A set of images is used to construct change sequences, where one image is presented repeatedly at regular times until a change occurs and a different image is presented for the remainder of the sequence. Each image is shown for 2 time steps and followed by 4 time steps of blank stimulation (zero input). This corresponds to a discretization of Δ*t* = 125 ms relative to the experimental task. During training, sequences are constructed from familiar images, while during testing, sequences are built from novel images.

#### Sequence construction

Each sequence consists of 8 image presentations in total, and there is a 4-time-step grace period of blank stimulation before the first image is presented. We allow 2 to 6 initial presentations before the change, creating 4 distinct “change times” for each ordered pair of images (***x***_*m*_, ***x***_*n*_). Thus, each ordered pair produces 4 sequences in which image ***x***_*m*_ changes to images ***x***_*n*_ at different times. In omission trials, we ensure that the omission does not coincide with the change time and that at least two initial presentations occur before the omission.

#### Image dataset

The images used to construct familiar and novel change sequences are from the TinyImageNet dataset ^49^, which is a reduced, 64 × 64 version of selected classes from the ImageNet dataset ^90^. We randomly selected 80 images from this dataset, spanning both the train and test splits. Those images were converted to grayscale using the ITU-R 601-2 luma transform, i.e.

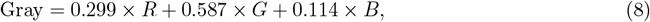

where *R, G*, and *B* are the red, green, and blue color channels. Example images are shown in Figure 1B.

Each grayscale image was first rescaled so that its pixel values are in the range [0, 1]. Then, for each pixel location (*k, k*^*′*^), we computed a pixel-wise mean ***µ***_*x*_(*k, k*^*′*^) and standard deviation ***σ***_*x*_(*k, k*^*′*^) across the entire dataset, and standardized every pixel by

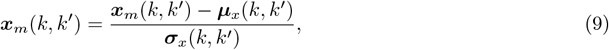

where ***x***_*m*_(*k, k*^*′*^) is the pixel value at row *k* and column *k*^*′*^ in image ***x***_*m*_. We then downsampled the standardized images to a resolution of 32 × 32 using average pooling with a 2 × 2 kernel and stride 2. Finally, the 2D images were reshaped (or flattened) into 1D vectors of 1024 features each, which were used to construct the change sequences. For every instance of training (i.e., random seed), two disjoint subsets (different for each instance) were randomly selected from this image dataset to be the familiar and novel sets.

### Detailed model architecture

Below, we give a more detailed description of the model architecture briefly outlined in the main text and depicted in Figure 2. Population sizes of the different types of neurons in the model are listed in Table S2.

#### Input layer (Layer 0)

The input layer simulates the retina in the visual pathway and is where the stimulus presentation occurs. At this layer, feedforward connections from the input stimulus project to the prediction error population in Layer 1 (*x*^err^) as well as the excitatory *z*, PV population, and SST (*θ*) populations in Layer 2.

#### Intermediate layer (Layer 1)

The intermediate layer corresponds to the intermediate stages of the visual processing pathway, such as the lateral geniculate nucleus (LGN) and other thalamic regions. It also captures the role of layer 4 in primary visual area, which is the main cortical recipient of thalamic input. At this layer, prediction error neurons *x*^err^ receive feedforward connections from the input layer as well as feedback from the excitatory *z* neurons in Layer 2, which computes a reconstruction, 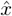, of the input stimulus, *x*. The feedforward connection to *x*^err^ is a fixed, identity mapping that directly maps *x* to *x*^err^ and participates in the computation of the prediction error at Layer 1, 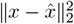. The feedback connections from Layer 2 are parameterized by a neural network 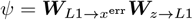. Therefore, Layer 1 also represents an intermediate step in the computation of the reconstruction 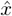 by receiving feedback connections ***W***_*z*→*L*1_ from Layer 2 and sending lateral connections 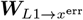 to local prediction error neurons. Biologically, regions in Layer 1, e.g., LGN, may also represent intermediate features of the raw stimulus before it is mapped to Layer 2. However, we choose not to explicitly model this step for simplicity.

#### Representation layer (Layer 2)

The sensory representation layer models early visual cortical areas– the primary (VISp) and secondary (VISl) areas of the mouse visual cortex, where experimental data was recorded ^23^. Functionally, it aligns with layer 2/3 in the laminar organization, which integrates input from earlier stages and participates in intracortical processing and feedforward projection to higher areas. There are four interneuron populations in this layer: excitatory, SST, VIP, and PV. The excitatory population consists of two subpopulations: representation (*z*) neurons and prediction error neurons (*z*^err^). The SST population comprises two subpopulations: SST (*θ*) and SST (*µ*^*p*^).

The excitatory *z* neurons receive feedforward connections from the input layer, parameterized by ***W***_*x*→*z*_. They also receive divisive inhibition through a fixed connection from the PV population as well as subtractive inhibition from the SST (*θ*) population through a connection parameterized by ***W***_*θ*→*z*_. Finally, the *z* population projects feedforward connections to Higher Areas A and B in the higher-level processing layer (Layer 3). Those connections are parameterized by 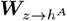 and 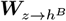.

The excitatory prediction error subpopulation, *z*^err^, receives fixed one-to-one connections from the *z* population, the SST (*µ*^*p*^) population, and the VIP population. These connections are all identity mappings and combine to compute the prediction error, 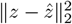, at Layer 2.

The VIP population primarily receives feedback connections from Higher Area B and the Reward area in the higher-level processing layer. The connections from Higher Area B are learned and parameterized by ***W***_*h*→vip_. The connection from the Reward area is fixed and acts as a mechanism for additive gain modulation.

The SST (*θ*) population receives inhibitory connections from the VIP population. These connections are parameterized by ***W***_vip→*θ*_, and their inhibitory effect is divisive, as shown in Eq. 16. SST (*θ*) also receives fixed feedforward connections from the input layer, parameterized by ***W***_*x*→*θ*_. These connections are used to compute a transient feedforward input, *θ*^ff^, which does not carry any stimulus-specific information; rather, it acts as a weak stimulus-triggered drive.

The SST (*µ*^*p*^) population receives only feedback connections from Higher Area A in the higher-level processing layer, which are parameterized by ***W***_*h*→sst_.

PV neurons receive feedforward connections from the input layer, parameterized by ***W***_*x*→pv_. These neurons also divisively inhibit the excitatory *z* neurons through a fixed one-to-one connection that maps a PV neuron’s response to its reciprocal, as shown in Eq. 14, 15, and 18.

#### Higher-level processing layer (Layer 3)

The higher-level processing layer represents the higher cortical areas involved in higher-order recurrent processing of input sequences and reward-based feedback modulation. There are three areas in this layer: Higher Area A, Higher Area B, and the Reward area. Each of these areas is recurrently connected with recurrent weight matrices 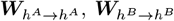, and ***W***_*v*→*v*_, respectively. Higher Areas A and B receive feedforward connections 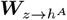 and 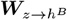 from the excitatory *z* population in Layer 2. The Reward area receives input from Higher Area B through connections parameterized by 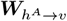 and sends a modulatory signal, *κ*, through a fixed feedback connection to the VIP population in Layer 2.

Since Higher Areas A and B integrate information from and send direct feedback to neurons in the representation layer, they are meant to correspond to higher visual areas in the mouse visual cortex, such as VISpm, VISrl, and VISam. Due to their recurrent connectivity and role in processing input sequences, we can also draw a correspondence between these areas and subregions in the prefrontal cortex (PFC) involved in working memory computation ^91^. The Reward area is involved in reward prediction and reward-guided decision making. Therefore, this area may correspond to regions in the mouse brain involved in reward-related behavior and reward-based feedback modulation, such as the orbitofrontal cortex (OFC), the dorsomedial striatum (DMS), and the ventral tagmental area (VTA) ^92–95^.

### Rationale behind model architecture

The architecture of the model is designed to reflect well-established principles of cortical circuit organization as it pertains to both experimental evidence and proposed biological implementations of hierarchical predictive coding.

PV neurons in Layer 2 receive primarily feedforward input from lower layers and no feedback input. This design is consistent with electrophysiological evidence that PV interneurons are strongly driven by incoming sensory signals and mediate feedforward inhibition of pyramidal cells without incorporating detailed top-down context ^11–13^. In contrast, VIP and SST neurons predominantly receive feedback from the higher areas, aligning with evidence that these cell types respond more robustly to top-down contextual and modulatory signals than direct sensory drive ^11–13,18,19^. The SST (*θ*) subpopulation is only assigned a weak, non-plastic feedforward input that produces a transient, non-tuned drive when a stimulus appears.

A hallmark of cortical microcircuitry incorporated in our model is the VIP → SST → Excitatory disinhibitory pathway. VIP neurons inhibiting SST cells and thereby relieving inhibition on excitatory neurons is a well-established motif in the cortex ^13,15^. Our model incorporates strong, plastic connections from VIP to SST (*θ*), consistent with evidence that VIP neurons preferentially target SST neurons, effectively silencing SST-mediated inhibition of excitatory cells ^10^. This mechanism is thought to underlie various forms of cortical gating ^14,40,96^.

The model incorporates a reward-based modulatory signal from the Reward area that specifically targets the VIP population. This design is grounded in extensive experimental evidence that VIP neurons are key targets of reinforcement learning signals and are strongly activated by reward or punishment^13,17,78,97–99^. In light of these findings, VIP neurons in our model are modulated by a feedback, additive gain signal based on expected reward, analogous to the uniform excitation of VIP cells observed experimentally under reinforcement.

The model distinguishes between two forms of inhibition onto Layer 2 excitatory neurons: PV neurons provide divisive inhibition while SST (*θ*) neurons provide subtractive inhibition. This design is rooted in the known functional differences between PV and SST cell impacts on pyramidal responses. PV cells target the perisomatic region of pyramidal neurons and tend to divide responses, while SST neurons target dendrites and tend to subtract from responses ^66^. The two forms of inhibition regulate the response gain and feature selectivity of excitatory neurons, respectively. These findings further support our assignment of distinct computational roles to different inhibitory subtypes.

The architecture is further informed by hierarchical predictive coding principles, wherein distinct populations encode predictions and prediction errors ^3,6,62,63^. Bastos et al. ^5^ proposed a canonical microcircuit for predictive coding where layer 2/3 pyramidal cells compare bottom-up input with top-down predictions, and inhibitory interneurons help compute these mismatches. In our model, we follow a similar scheme. For example, excitatory *z* neurons represent inferred latent states of sensory observations, while *z*^err^ neurons compute the mismatch between inferred and expected states, which are encoded by inhibitory neurons receiving feedback from higher areas. However, in contrast to Bastos et al. ^5^, our model provides a specific assignment of computational roles to different interneuron subtypes in a manner that is consistent with their observed connectivity and functional properties. For example, in our model, the SST (*µ*^*p*^) population explicitly encodes top-down predictions, while the VIP population encodes the uncertainty associated with those predictions.

### Algorithmic Implementation

We model the posterior distribution given an input stimulus, *q*(*z* | *x*), and the prior distribution given sequence history, *p*(*z* |ℋ ), as Gaussian distributions. The posterior has mean ***µ***^*q*^ and diagonal covariance diag(***σ***^*q*^), while the prior has mean ***µ***^*p*^ and diagonal covariance diag(***σ***^*p*^).

At time step *t*, connections from Layer 3 to Layer 2 compute the parameters of the prior distribution as follows.

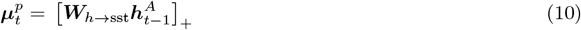

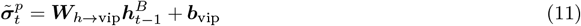

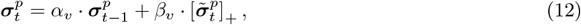

where *α*_*v*_ and *β*_*v*_ are scalar gain hyperparameters that balance the contributions of previous values (history) and the current input, [·]_+_ is the rectified linear unit, and ***b***_vip_ is a fixed bias initialized randomly before training. Note that there is a 1-time-step delay in the feedback projections from the higher areas in Layer 3 to the interneuron populations in Layer 2.

The parameters of the posterior distribution are computed by feedforward input to the excitatory and PV neurons as follows. Note that there is a 1-time-step delay from the stimulus in the input layer to the neurons in Layer 1 and Layer 2.

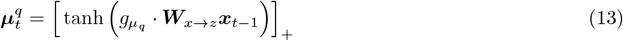

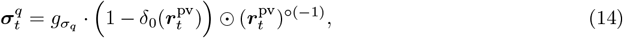

where δ_0_ is the Dirac Delta function at 0, ⊙ is the elementwise (or Hadamard) product, (·)^°(−1)^ is the elementwise (or Hadamard) inverse, 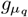 and 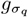 are fixed gain parameters, and 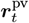 is the PV population response, computed as

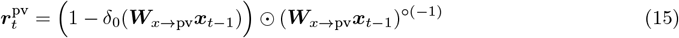

Note that this formulation allows ***σ***^*q*^ to contain zero elements, which violates the condition that *σ >* 0 for a Gaussian distribution. However, this only occurs during blank stimulation when there is zero input. In this case, there is no feedforward drive to the excitatory and PV neurons and the posterior distribution is a Dirac Delta function at **0**.

The activities of the SST *θ* subpopulation are computed as

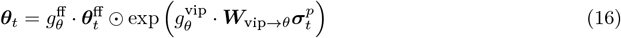

where ***θ***_ff_ is the feedforward input to the *θ* population, computed as

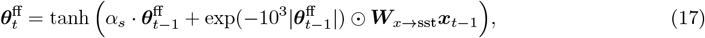

and 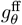 and 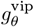 are fixed gain parameter controlling the magnitude of feedforward input and VIP inhibition, respectively. Note that since ***W***_vip→sst_ consists of only non-positive weights, the function divisive inhibition. exp 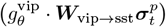 is a non-increasing function of 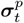. Therefore, the effect of 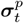 on SST neurons is divisive inhibition.

To compute the activities of the excitatory *z* subpopulation, we first combine the feedforward input ***µ***^*q*^ with from ***r***^pv^ to compute a net feedforward drive as follows.

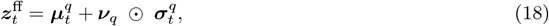

where ***ν***_*q*_ ∼ 𝒩 (**0, *I***). Note that 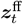 corresponds to a sample from the posterior distribution.

Then, we compute the inhibitory input from the SST *θ* neurons to the excitatory *z* neurons as

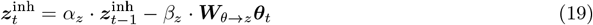

The final activations of the *z* neurons are then computed as

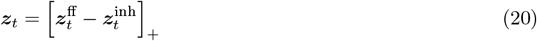

The activities of Higher Area *A* and Higher Area *B* are updated through recurrent connections and feedforward input from the excitatory *z* neurons as follows.

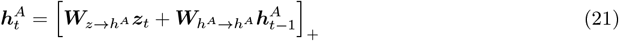

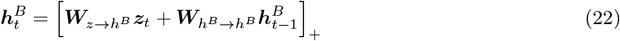

The Reward area in the higher-level processing layer receives feedforward input only from Higher Area B. Its activities are computed as

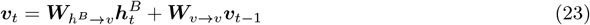

The decision-making component of the model now proceeds as follows. First, the response average of half the neurons in the Reward area is taken to represent the value of licking, while the average of the other half represents the value of not licking. More precisely, those values are computed as follows.

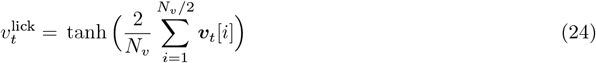

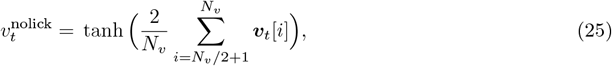

where *N*_*v*_ is the number of neurons in the Reward area.

The probability of licking is calculated as

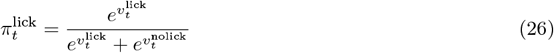

The action at time *t* is then decided via an *ϵ*−greedy exploration strategy as follows.

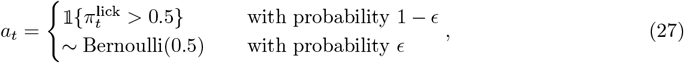

where *ϵ* ∈ [0, 1] is the exploration probability.

The predicted value of the chosen action is calculated as

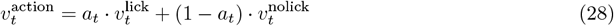

This predicted value is used to calculate a feedback signal that acts as a modulatory gain on the activities of VIP neurons (represented by ***σ***^*p*^). This gain is calculated as

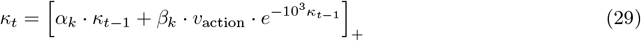

The VIP response (***σ***^*p*^, Eq. 12) is then updated as follows:

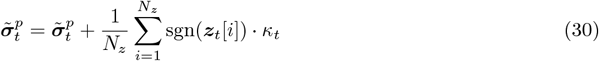

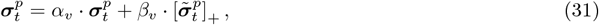

where *N*_*z*_ is the number of excitatory (*z*) neurons. We make the final modulatory gain signal dependent on the level of excitatory activity by considering fixed *binary* connections (computed by the sgn function) from the excitatory (*z*) neurons onto the VIP neurons.

Top-down predictions from the higher-level processing and the representation layers are now computed as follows.

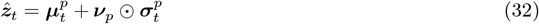

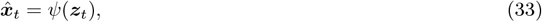

where ***ν***_*p*_∼ 𝒩 (**0, *I***), and *ψ* is the neural network parameterizing the feedback mapping from the representation layer to the input layer (see section detailed model architecture above). Note that 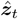 is the higher area’s prediction of the excitatory *z* activities, and 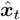 is the representation layer’s prediction (or reconstruction) of the input image.

The activities of the prediction error neurons at the representation and the input layers are computed as

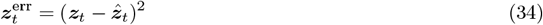

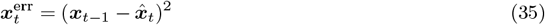

Finally, after executing action *a*_*t*_ and receiving reward *R*_*t*_, the model computed a reward prediction error as follows.

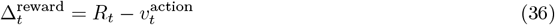

For initialization at *t* = 0, we set 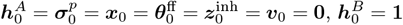, and *κ*_0_ = 0.

#### Mapping algorithmic nodes to interneuron populations

##### PV neurons

PV neurons receive feedforward drive from the input layer and represent the precision of the posterior distribution *q*(***z*** | ***x***), or the reciprocal of the standard deviation,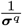. This aligns with their role in divisively inhibiting the excitatory *z* population when sampling from the posterior distribution in Eq. 18.

##### VIP neurons

VIP neurons represent the uncertainty of top-down predictions coming from the higher-level processing layer. More precisely, their activities correspond to the standard deviation ***σ***^*p*^ of the prior distribution. This aligns with their role in subtractively inhibiting the excitatory prediction error neurons when computing the MSE between actual and predicted *z* activities, as shown in Eq. 32 and 34.

##### SST neurons

There are two subpopulations within the SST population in our model: SST (*µ*^*p*^) and SST (*θ*). The SST *µ*^*p*^ population represents the mean of the prior distribution. Similar to VIP neurons, they participate in sampling a prediction 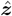 from the prior distribution, which is used to compute the prediction error at the representation layer by inhibiting the excitatory prediction error neurons (as shown in Eq. 32 and 34).

The SST (*θ*) population receives inhibitiory input from the VIP neurons and weak feedforward drive from the input layer. This population serves as part of the VIP-SST disinhibitory pathway and acts as the primary source of inhibition to the excitatory neurons.

##### Excitatory neurons

The excitatory population in our model consists of two subpopulations: representation neurons (*z*) and prediction error neurons (*z*^err^). Representation neurons receive feedforward drive from the input layer, representing the mean of the posterior distribution ***µ***^*q*^. This feedforward drive, combined with divisive inhibition from PV neurons, results in a net feedforward input 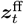 which corresponds to a sample from the posterior distribution as shown in Eq. 18. Since these representation neurons are inhibited by SST neurons, their final activities represent a posterior latent state modified by inhibitory input.

The prediction error neurons *z*^err^ receive excitatory input from the *z* neurons as well as inhibitory input from the SST (*µ*^*p*^) subpopulation and the VIP population. Therefore, these neurons represent the prediction error between the top-down prediction and the bottom-up input at the representation layer, as shown in Eq. 34.

### Training

#### Objective Functions

In line with the theory of hierarchical predictive coding theory ^3^, we train the model to minimize the activities of the prediction error neurons in the input and representation layers. In addition to these two objectives, the model is trained to optimize energy efficiency and behavioral performance. Energy efficiency is represented by the *l*_1_ − norm of the excitatory representation units, *z*, while behavioral performance is optimized by minimizing the reward prediction error. More precisely, for a given sequence *d* of length *T*, the prediction error loss functions are given by Eq. 3, while the energy efficiency and reinforcement learning objectives are given by Eq. 5 and 4, respectively. The model parameters are trained with backpropagation to minimize the final objective function given by Eq. 6.

Note that our behavioral training approach corresponds to an actor-critic setup in the Reinforcement Learning (RL) literature ^100^. Specifically, the Reward area in our model acts as the critic by predicting the values of different actions. The model then chooses an action based on the predicted values, and the reward prediction error is used to improve the model’s predictions.

#### Annealing Schedules

Each of the scalar weights in Eq. 6, except for *λ*_1_, is annealed throughout training according to an annealing schedule that guides the optimization of different objectives. For *y* ∈ {2, energy, action }, those annealing schedules, as a function of training epoch *j*, are

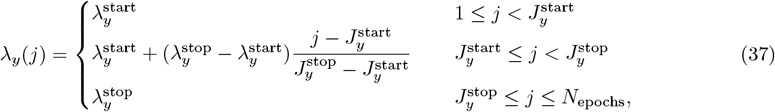

where *N*_epochs_ is the total number of training epochs. In words, this annealing schedule holds *λ*_*y*_ constant at 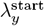 from the first epoch up to epoch 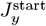. Between epoch 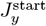 to epoch 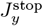, *λ*_*y*_ ramps up linearly from 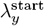 to 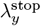. From epoch 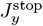 onward, *λ*_*y*_ is held constant at 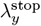.

The exploration probability *ϵ* is also annealed throughout training according to Eq. 37; albeit with parameters *ϵ*^start^ and *ϵ*^stop^ indicating initial and final values. In this case, however, *ϵ*^start^ *> ϵ*^stop^, which results in *ϵ* linearly decreasing between epochs 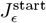 and 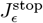 during training.

The annealing parameters 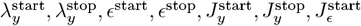 and 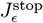 are all hyperparameters whose values are adjusted to balance the performance of the model and the replication of experimental observations (see Table S1).

#### Learned and Fixed Parameters

To simplify the model as much as possible, some connections are learned during training and some are fixed (Figure 2). The learned connections are parameterized by weight matrices that are adjusted with backpropagation by optimizing the loss function in Eq. 6 (see the above section on model architecture). All learned connections are affected by all losses; however, long-range, inter-area feedforward connections progressively become fixed at the later stages of training. That is, when training the model for 70 epochs, feedforward connections from the input layer become fixed at epoch 25, and feedforward connections from Layer 2 become fixed at epoch 35. Since 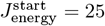, this means that feedforward connections from the input layer are mainly affected by the gradients of the prediction error losses, 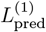 and 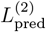, and the behavioral objective, *L*_action_.

Fixed connections are not parameterized by weight matrices; rather, they are determined by their role in the algorithmic implementation outlined above. The only exception is the feedforward connection from the input layer to the SST *θ* subpopulation, which is parameterized by ***W***_*x*→sst_ and is initialized and fixed before training. How the fixed connections are determined by algorithmic roles is described below.

#### Enforcing Dale’s law

To maintain biological plausibility, all connections in our model are constrained to obey Dale’s law, including both learned and fixed connections. That is, connections originating from excitatory and inhibitory neurons are parameterized by nonnegative and nonpositive weights, respectively. The only exception to this constraint are inter-layer connections, which are allowed to be heteregenous, i.e. include either positive or negative weights.

For learned connections, Dale’s law is enforced by constraining the connection weight matrix ***W*** to be either nonnegative or nonpositive depending on the presynaptic population. The only intra-layer learned connections in our model are those from VIP to SST (*θ*), parameterized by ***W***_vip→*θ*_, and those from SST (*θ*) to the excitatory *z* neurons, parameterized by ***W***_*θ*→*z*_. Both connections are inhibitory and their weights are forced be negative through rectification as follows.

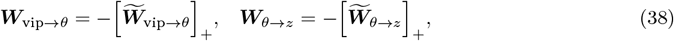

where 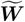 is the unconstrained weight matrix for each connection.

For fixed *intra-layer* connections, their role in the algorithmic implementation of the model determines whether they are inhibitory or excitatory. Since these connections are not parameterized, their algorithmic roles were specifically chosen to maintain consistency with Dale’s law. Below is a description of how each of these connections obeys Dale’s law based on algorithmic role.

- <u>Connections to the excitatory prediction error neurons</u> Excitatory prediction error neurons, *z*^err^, represent the prediction error at Layer 2. Their activities are computed according to Eq. 34, which can be re-written as

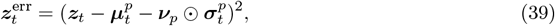

where ***ν***_*p*_∼ 𝒩 (**0, *I***). This computation involves excitatory connections from the *z* population and inhibitory connections from the SST (*µ*^*p*^) and VIP neurons. To see how these connections obey Dale’s law, we can think of them as being parameterized by fixed weight matrices 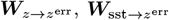, and 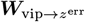, respectively, such that

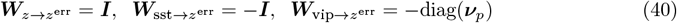

The activities of *z*^err^ can then be expressed as

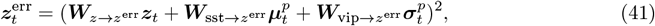

where 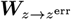 contains only nonnegative weights and 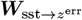 contains only nonpositive weights. However, since ***ν***_*p*_ is drawn from a standard normal distribution, 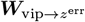 can contain positive weights at entries where ***ν***_*p*_ is negative. This issue can be resolved by noting that VIP neurons can have a net excitatory effect on the *z*^err^ neurons through the VIP-SST disinhibitory pathway. That is, we can write the top-down prediction 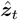 in Eq. 32 equivalently as

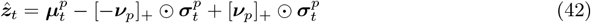

This gives us the following alternative implementation for computing the activities of *z*^err^.

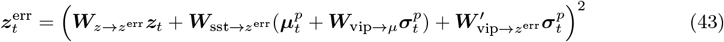

where

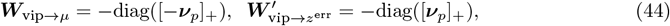

and 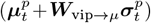 is the SST (*µ*^*p*^) response in this alternative implementation. This is equivalent to our original implementation but maintains strict nonpositivity for all connections originating from VIP neurons. For simplicity, however, we choose to absorb ***W***_vip→*µ*_ and 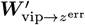 into one weight matrix 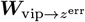.
- <u>Connections from PV neurons to excitatory *z* neurons</u> This connection comes in when computing the net feedforward input to the excitatory *z* neurons in Eq. 18. PV neurons are assumed to exert *divisive* inhibition. Therefore, in our implementation, they are computed as the inverse of the standard deviation 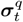, as shown in Eq. 14 and 15. Note that, since ***ν***_*q*_ is drawn from a standard normal distribution and can contain negative entries, we run into an issue similar to the one described above. However, we can again get around this issue by using an alternative implementation that utilizes the disinihbitory pathway between PV, SST (*θ*), and excitatory *z* neurons.

#### Parameter Initialization

All weight matrices, except for ***W***_*x*→sst_ and ***W***_*θ*→*z*_, were initialized using Xavier initialization ^101^. For a given weight matrix ***W*** ∈ ℝ^*m×n*^, its values were initialized as

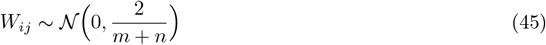

The values of ***W***_*x*→sst_ were initialized from a normal distribution with mean 1 and standard deviation 10^−3^, while those of ***W***_*θ*→*z*_ were initialized from a normal distribution with mean 0.1 and standard deviation 10^−3^. Values of the bias vector ***b***_vip_ were drawn from a normal distribution with mean 0.8 and standard deviation 0.3.

The weight matrix 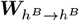, which parametrizes the recurrent connections in Higher Area B, was constrained throughout training to have a maximum norm of 0.5 to ensure stable dynamics. Although Higher Area A and the Reward area also have recurrent connections, leaving their connectivity matrix unconstrained did not result in unstable dynamics during training. Therefore, we chose not to constrain the connectivity of those areas.

### Response adaptation model with Hebbian learning

The response adaptation model is a firing rate model that consists of three interneuron populations: excitatory (exc), VIP, and SST. The connectivity between these populations, shown in Figure 8C, was designed to match the connectivity between the same populations in our original model. The task setup and task parameters used for this model were also the same as the original model; however, we used a finer discretization of Δ*t* = 50 ms for constructing image sequences, corresponding to 5 time steps for image presentations and 10 time steps for blank screens.

To differentiate from notation used in our original model description, we will use ***C*** to refer to connectivity weight matrices. The firing rate dynamics for each population *y* (*y* ∈ { exc, vip, sst } ) are described by

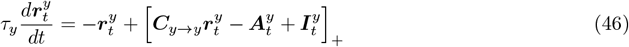

where:

- 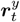 is the firing rate of population *y* at time *t*.
- *τ*_*y*_ is the membrane time constant for neurons in population *y*.
- ***C***_*y*→*y*_ is the synaptic weight matrix for recurrent connections in population *y*.
- 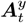 is the vector of adaptation variables for neurons in population *y*.
- 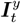 is the external input to population *y*, computed for each population based on the model’s connectivity as follows.

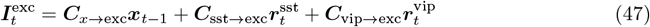

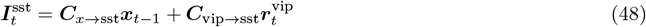

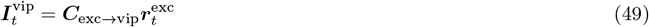

where ***x***_*t*−1_ is the input image at time *t* − 1. Note that, similar to our original model, we implement a one-time-step delay between the input and the interneuron populations.

The adaptation variables in 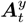 represent the adaptation effects in neurons of population *y* and evolves according to:

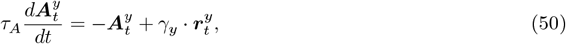

where *τ*_*A*_ is the adaptation time constant (same for all populations) and *γ*_*y*_ is the adaptation strength for neurons in population *y*. All responses ***r*** and adaptation variables ***A*** were initialized to **0** at time step *t* = 0 in the sequence.

When conducting simulations with Hebbian learning, synaptic weights *C*_*ij*_ were updated using a Hebbian learning rule:

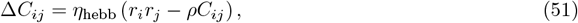

where *η*_hebb_ is the learning rate, *r*_*i*_ and *r*_*j*_ are the post- and presynaptic firing rates, respectively, and *ρ* is a weight decay factor. At every time step in the sequence, this update was performed for all weight matrices, except for the feedforward connections to SST neurons (***C***_*x*→sst_) which we treat as fixed, similar to our original model.

All connectivity weight matrices were constrained to obey Dale’s law by rectifying weights in randomly-initialized unconstrained weight matrices 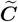 as follows.

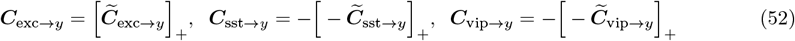

When using Hebbian learning, those constraints were re-enforced after every Hebbian update step. To prevent unlimited growth, synaptic weight matrices were normalized such that the vector of incoming weights onto each neuron *i* had unit norm. That is, each weight *C*_*ij*_ from neuron *j* to neuron *i* was normalized as

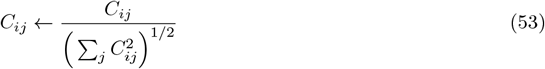

Unconstrained matrices 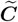 were initialized using the same method in Eq. 45.

All hyperparameters in this model were chosen to best replicate the experimental novelty effects in Garrett et al. ^23^, and are shown in Table S3.

## 5 Data Availability

All the experimental data analyzed in this manuscript is part of the existing, publicly available Allen Brain Map – Visual Behavior (2P) dataset introduced in Garrett et al. ^23^. The pre-processed calcium imaging data used to generate the experimental figures is available for download at https://doi.org/10.6084/m9.figshare.29207546.v2. The corresponding raw data is available for download in Neurodata Without Borders (NWB) format via the AllenSDK at https://portal.brain-map.org/circuits-behavior/visual-behavior-2p.

## 6 Code Availability

The code and data to reproduce the results in this article are available at https://github.com/HChoiLab/cortical_circuit_model_for_novelty_encoding/.

## 7 Acknowledgment

This work was supported by a Alfred P. Sloan Research Fellowship in Neuroscience to H.C. We thank Marina Garrett for insightful feedback on the manuscript and guidance through the Allen Institute’s Visual Behavior-2P dataset. We also thank Stefan Mihalas and Eric Shea-Brown for helpful discussions.

## 8 Author Contributions

Conceptualization: A.S. and H.C.; Methodology: A.S. and H.C.; Software, visualization, and formal analysis: A.S.; Investigation and Validation: A.S. and H.C.; Writing-original: A.S.; Writing-editing: A.S. and H.C.; Funding acquisition, Project administration, and Supervision: H.C.

## 10 Supplementary Information

### Supplementary Tables

**Table S1:**
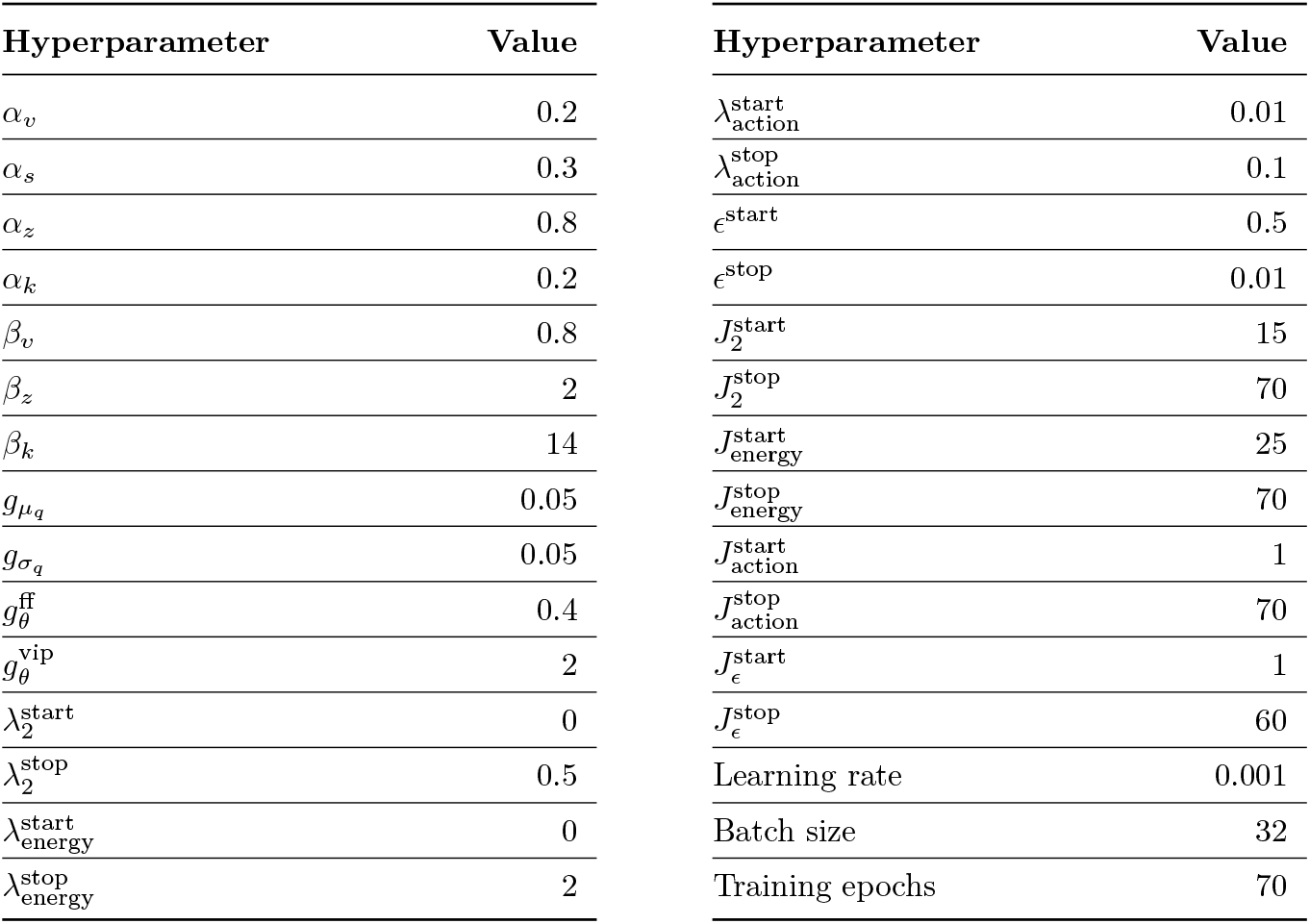
Hyperparameters for the proposed model

**Table S2:** Population sizes in the proposed model

| Population | Number of Neurons |
| --- | --- |
| Layer 1 prediction error ($x^{\text{err}}$) | 1024 |
| Excitatory representation ($z$) | 64 |
| Excitatory prediction error ($z^{\text{err}}$) | 64 |
| PV | 64 |
| VIP | 64 |
| SST ($\theta$) | 64 |
| SST ($\mu^p$) | 64 |
| Higher Area A | 128 |
| Higher Area B | 128 |
| Reward Area | 128 |

**Table S3:** Hyperparameters for response adaptation model with Hebbian learning

| Hyperparameter | Value |
| --- | --- |
| $\tau_{\text{exc}}$ | 0.3 |
| $\tau_{\text{vip}}$ | 0.3 |
| $\tau_{\text{sst}}$ | 0.3 |
| $\tau_A$ | 1.0 |
| $\gamma_{\text{exc}}$ | 20 |
| $\gamma_{\text{vip}}$ | 20 |
| $\gamma_{\text{sst}}$ | 10 |
| $\eta_{\text{hebb}}$ | $10^{-5}$ |
| $\rho$ | 0.001 |
| Simulation $dt$ | 0.1 |

### Supplementary Figures

**Figure S1:**
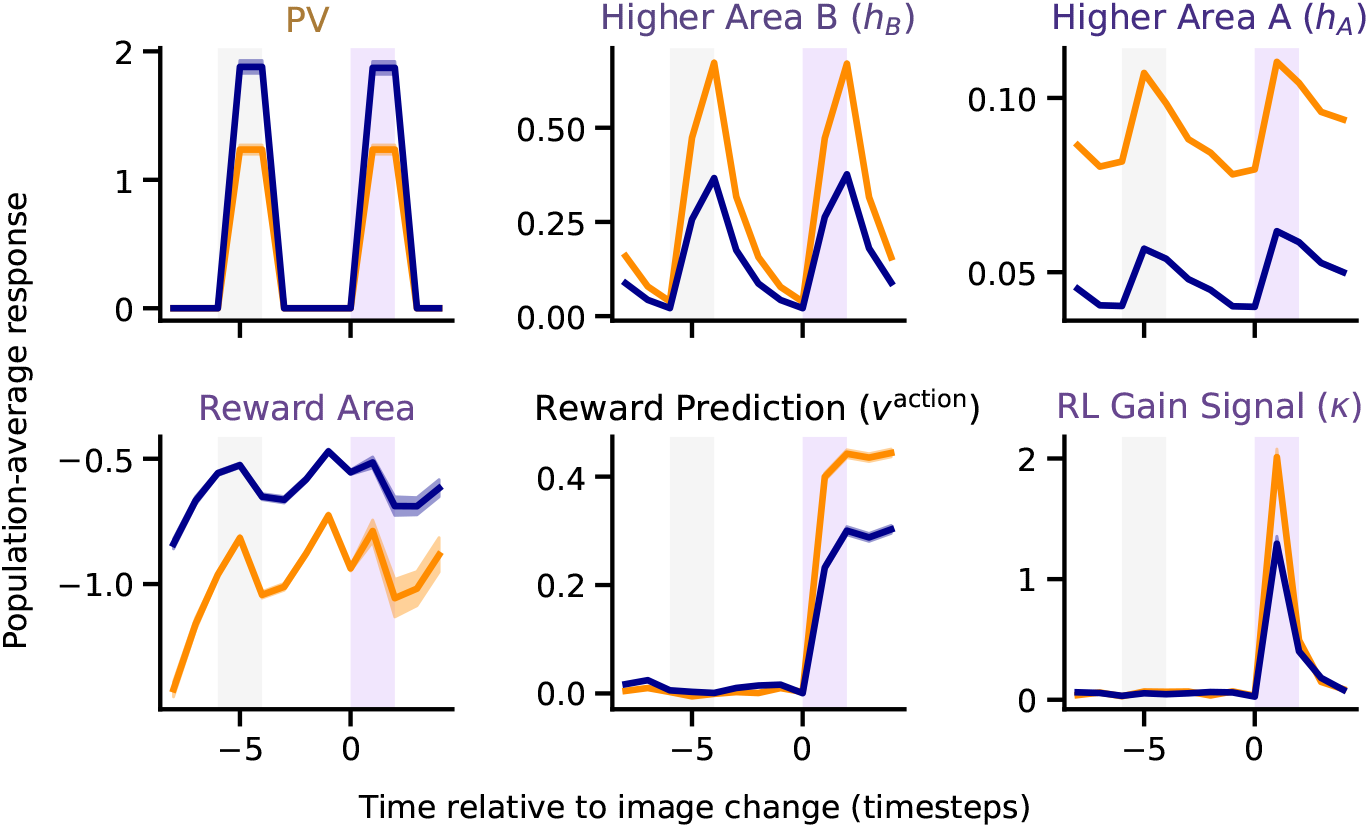
Full model change responses. Change responses in the full model (perception + action) of populations not included in the main text. The PV responses correspond to the variable ***r***^pv^ in the algorithmic implementation (see Methods). Responses shown are averages across neurons, trials, and training instances (n = 16 random seeds). Shaded error bars represent SEM.

**Figure S2:**
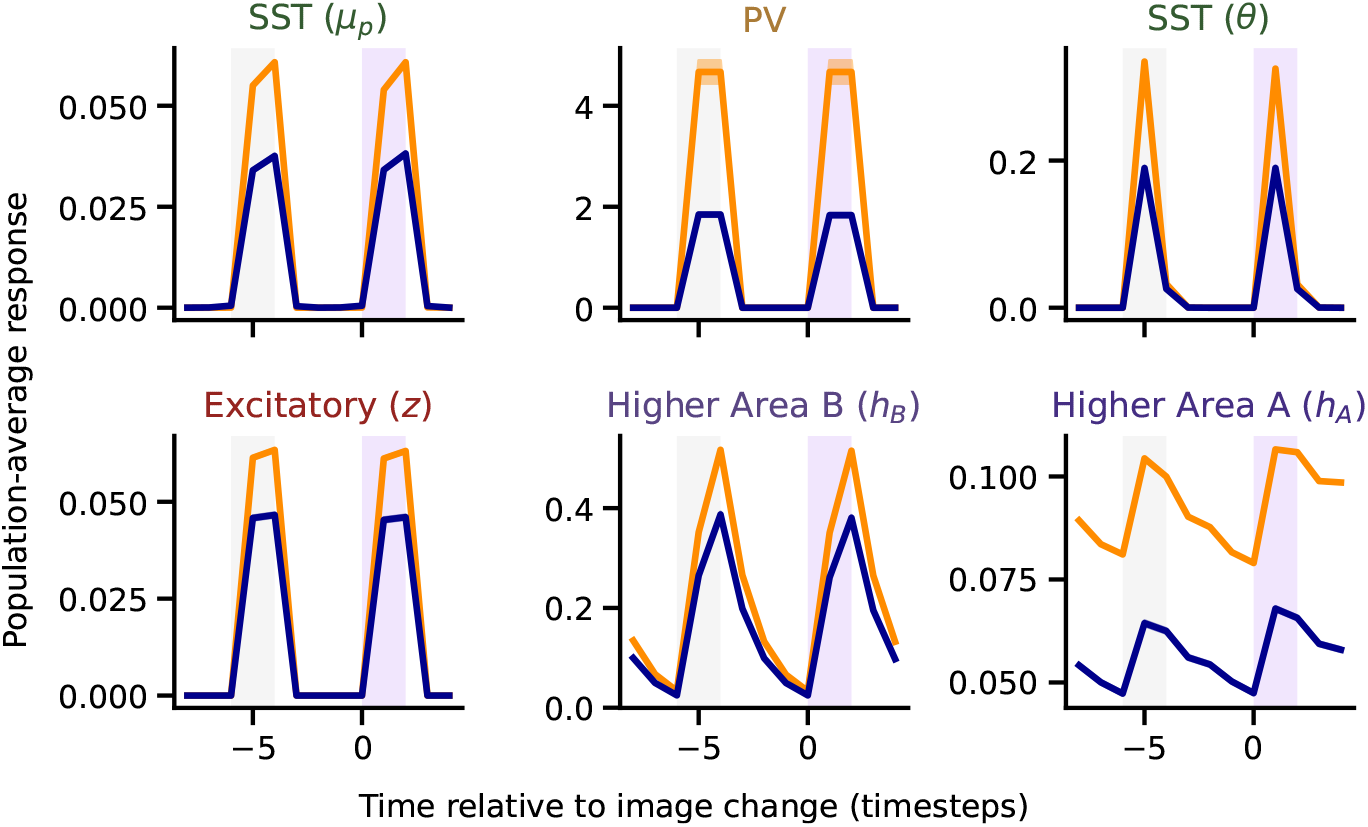
Change responses of other populations without reinforcement learning. Change responses of populations not included in the main text for the perception-only model (i.e, without reinforcement learning). These responses are generated with the same model setup as Figure 6A. Responses are averages across neurons, trials, and training instances (n = 16 random seeds). Shaded error bars represent SEM.

**Figure S3:**
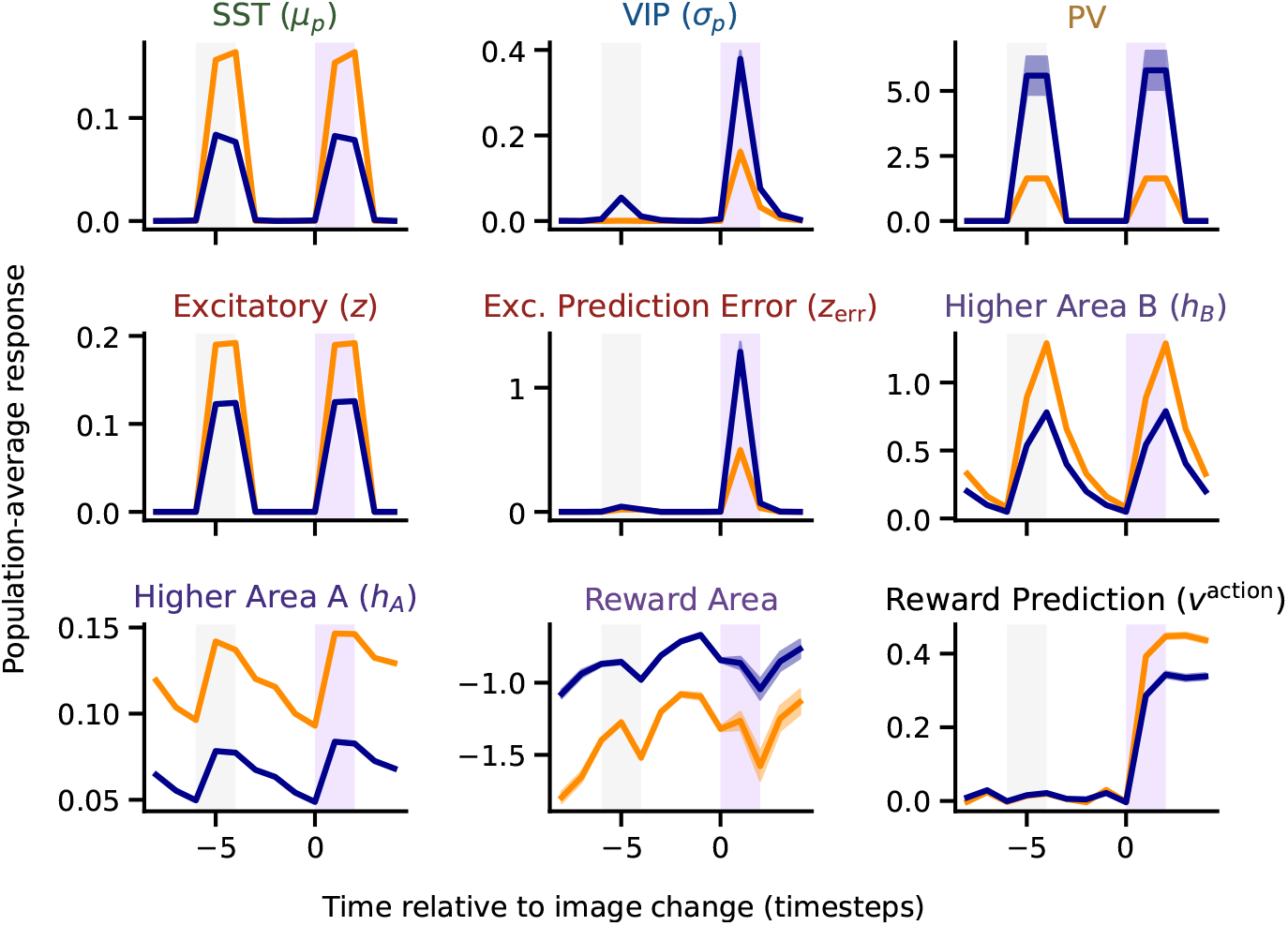
Change responses of other populations without energy efficiency. Change responses of populations not included in the main text for the model without energy efficiency optimization. These responses are generated with the same model setup as Figure 6B. Responses are averages across neurons, trials, and training instances (n = 16 random seeds). Shaded error bars represent SEM.

**Figure S4:**
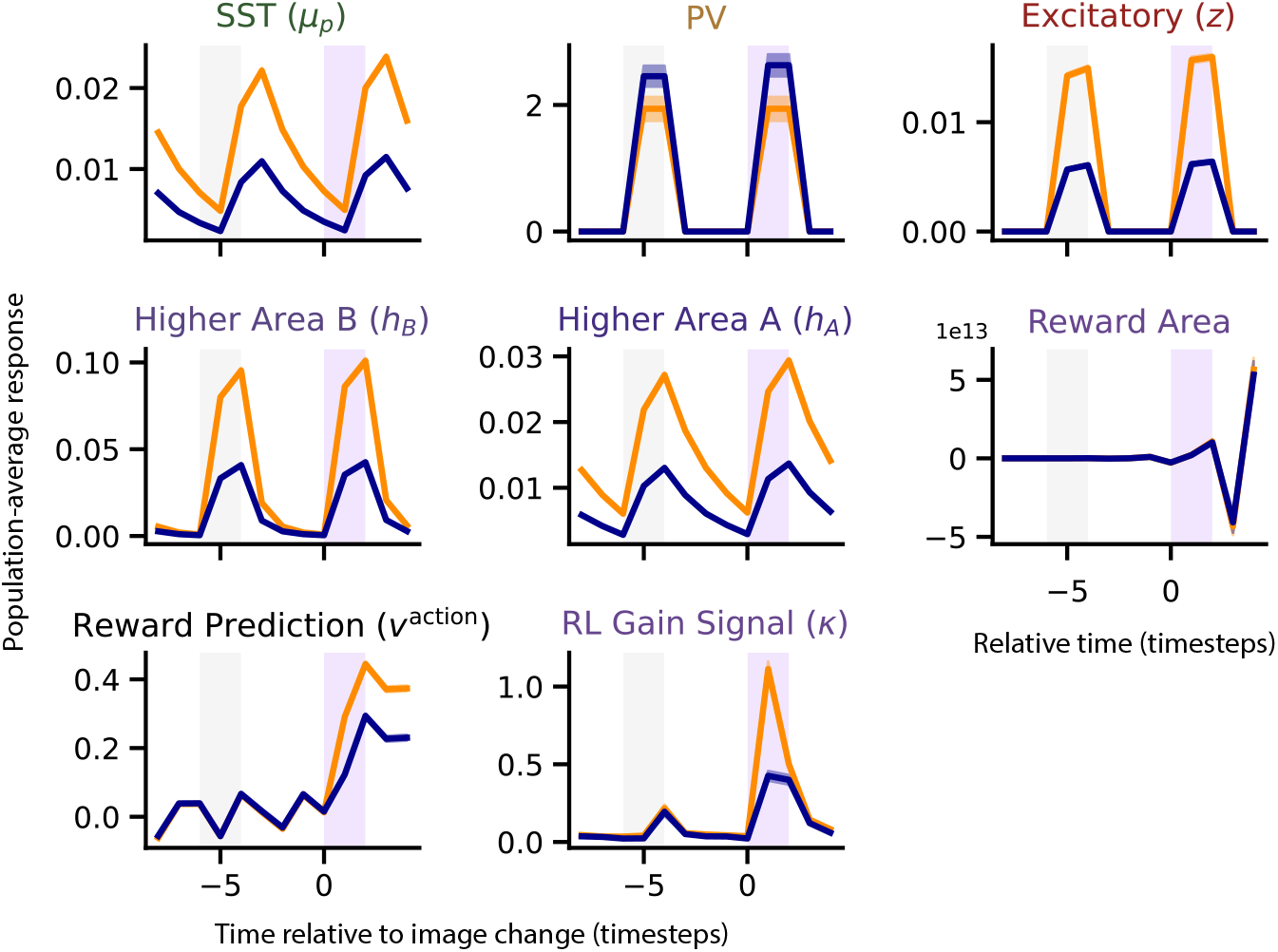
Change responses of other populations without predictive coding. Change responses of populations not included in the main text for the model without prediction error minimization. These responses are generated with the same model setup as Figure 6C. Responses are averages across neurons, trials, and training instances (n = 16 random seeds). Shaded error bars represent SEM.

**Figure S5:**
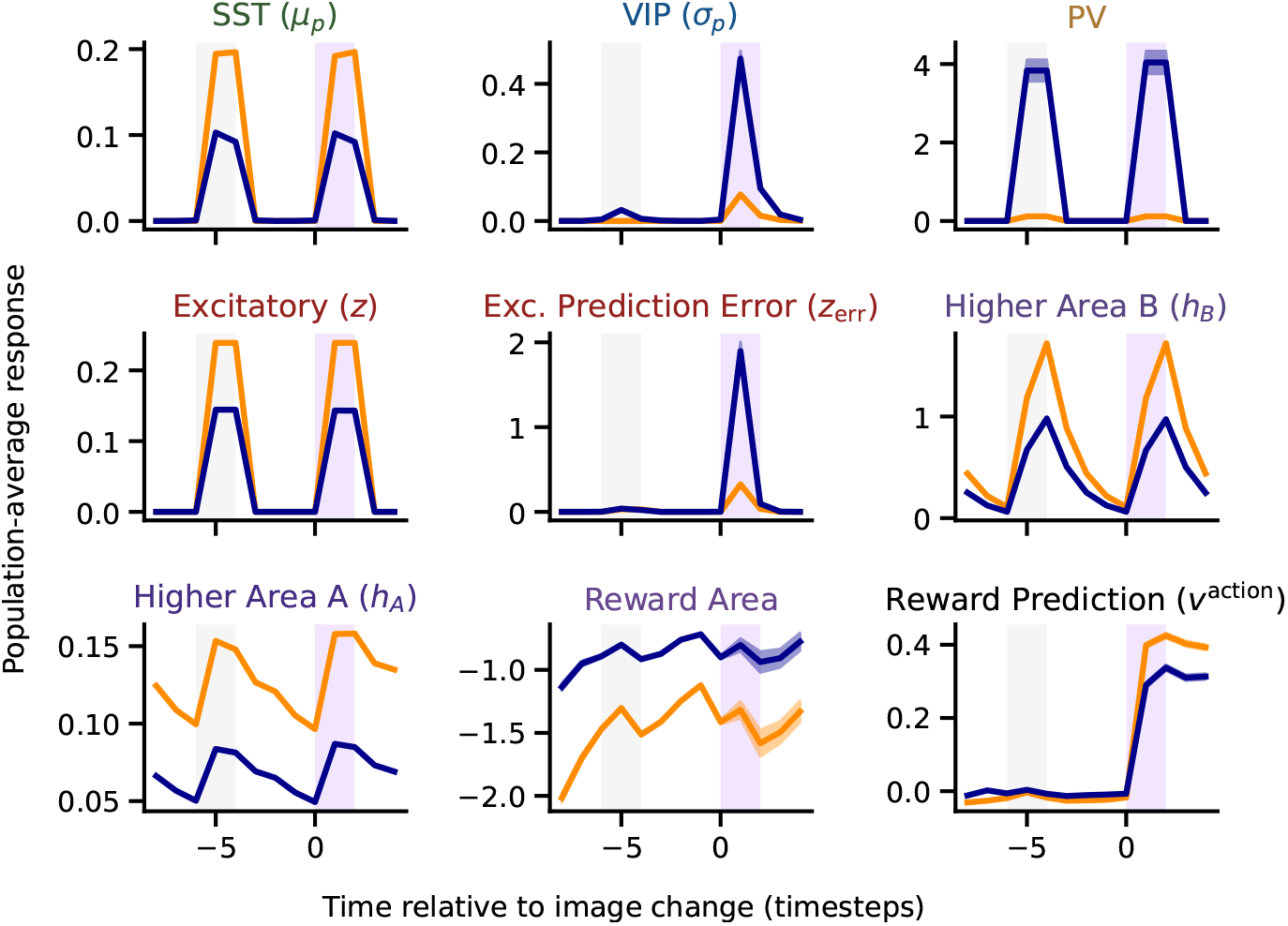
Change responses of other populations without SST-Exc inhibition. Change responses of populations not included in the main text for the model with the connection between SST (*θ*) and Excitatory (*z*) neurons removed. These responses are generated with the same model setup as Figure 7A. Responses are averages across neurons, trials, and training instances (n = 16 random seeds). Shaded error bars represent SEM.

**Figure S6:**
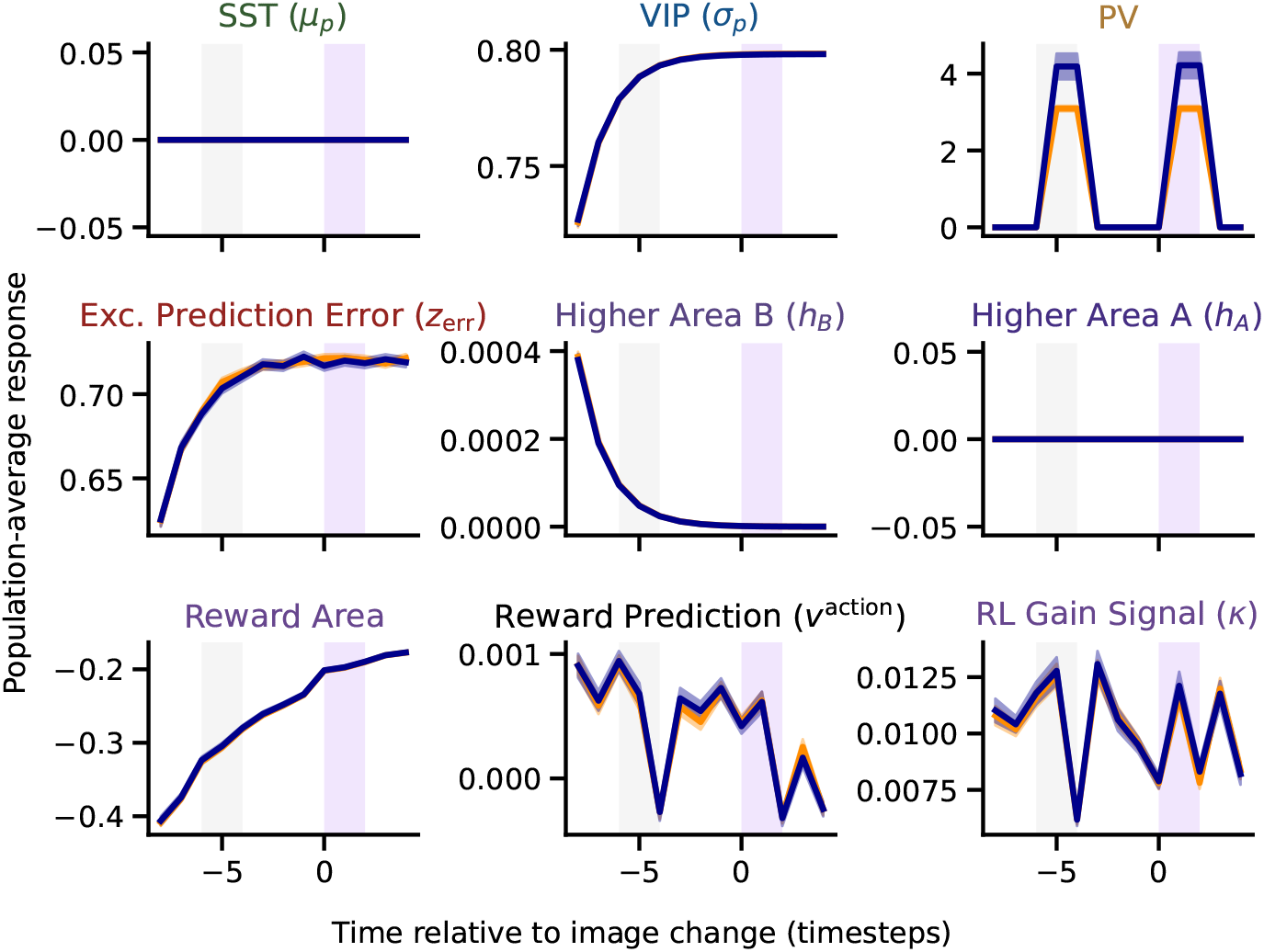
Change responses of other populations without VIP-SST disinhibition. Change responses of populations not included in the main text for the model with the connection between VIP and SST (*θ*) neurons neurons removed. These responses are generated with the same model setup as Figure 7E. Responses are averages across neurons, trials, and training instances (n = 16 random seeds). Shaded error bars represent SEM.

